# CDK4 deletion in mice prevents fat accumulation and increases endurance capacity through activation of estrogen-related receptor (ERR)-driven oxidative metabolism in skeletal muscle

**DOI:** 10.1101/2022.03.03.482783

**Authors:** Valentin Barquissau, Nadège Zanou, Sarah Geller, Judit Castillo-Armengol, Flavia Marzetta, Katharina Huber, Dorian Ziegler, Isabel Lopez-Mejia, Joan Blanco Fernandez, Catherine Roger, Nicolas Guex, Frédéric Preitner, Jean-Marc Vanacker, Lluis Fajas

## Abstract

Cyclin-dependent kinase 4 (CDK4) canonical role is to control cell cycle progression from G1 to S phases. However, recent studies reported that CDK4 regulates energy metabolism in non-proliferating cells such as hepatocytes or adipocytes. The objective of our work is to study CDK4 function in skeletal muscle using a model of mice lacking CDK4 (*cdk4*^*-/-*^). By coupling treadmill running to indirect calorimetry, we show that *cdk4*^*-/-*^ mice display improved endurance and higher capacity to use fat as fuel during exercise. Isolated muscles lacking CDK4 are more resistant to fatigue in response to repeated contractions and have increased oxidative capacity and mitochondrial content compared to *cdk4*^*+/+*^ muscles. Transcriptomic analysis reveals upregulation of genes controlled by the nuclear receptors estrogen-related receptors (ERRs) in *cdk4*^*-/-*^ skeletal muscle, associated with elevated levels of the ERR co-activator PGC1a. Supporting in vivo results, C2C12 myotubes treated with a CDK4 inhibitor have increased mitochondrial oxygen consumption, PGC1α expression and ERR transcriptional activity measured by a luciferase reporter. In normal housing conditions, *cdk4*^*-/-*^ mice show an increased basal metabolic rate and are resistant to weight gain and fat accumulation. In conclusion, our study uncovers a role for CDK4 in the control of skeletal muscle metabolism. Moreover, CDK4 inhibition may be an alternative strategy against obesity-associated metabolic disorders.

## Introduction

Cell cycle regulators such as cyclins, cyclin-dependent kinases (CDKs) and E2F transcription factors control the sequential progression through the cell cycle by coordinating the expression of genes required to complete each phase. During the G1 phase, CDK4 and CDK6, after phosphorylation by CDK-activating kinases (CAK), become activated through binding of mitogen-induced D-type cyclins (Malumbres, 2014). The newly formed cyclinD-CDK4/6 complexes, in turn, phosphorylate the retinoblastoma protein (RB) and relieve the E2F-mediated repression of the expression of genes involved in the G1-to-S transition enabling the entry into the S phase. Consistent with a role in cell proliferation, CDK4/6 expression and activity were found to be deregulated in several types of cancer (Malumbres & Barbacid, 2009). This led to the development of CDK4/6 inhibitors, which are currently used for the treatment of certain types of breast cancers (George *et al*, 2021)).

Beyond the canonical role in the control of cell cycle and cell proliferation, studies using murine models of CDK4 deficiency uncovered novel functions for this kinase in non-dividing cells. Mice lacking CDK4 are viable but have reduced size and impaired fertility. However, isolated fibroblasts lacking CDK4 proliferate normally, suggesting that redundant mechanisms compensate for the lack of CDK activity during the cell cycle progression (Rane *et al*, 1999) (Malumbres *et al*, 2004). Nevertheless, *cdk4* full-knockout mice show an early type 1 diabetic phenotype consecutive to a reduction in pancreatic beta cell number, leading to death around 6 months of age (Rane *et al*., 1999). When the expression of CDK4 was rescued specifically in the insulin-producing cells of the *cdk4* full-knockout mice, the beta cell number, insulin production, blood glucose and lifespan were completely normalized ((Martin *et al*, 2003); in our study from here onwards, these mice are referred to as *cdk4*^*-/-*^ mice, compared to their control littermates *cdk4*^+/+^). However, despite the restoration of pancreatic beta cell development and function, *cdk4*^*-/-*^ mice still display whole-body and tissue-specific metabolic alterations. C*dk4*^*-/-*^ mice have a lower body weight (Martin *et al*., 2003) and decreased fat accumulation in white adipose tissue (Lagarrigue *et al*, 2016). Moreover, *cdk4*^*-/-*^ are resistant to fatigue when submitted to a running test (Lopez-Mejia *et al*, 2017). Indeed, it was demonstrated in this study that CDK4 phosphorylates AMP-activated protein kinase (AMPK) on serine 377 to inhibit its activity in mouse embryonic fibroblasts (MEF), thereby inhibiting fatty acid oxidation; consequently, *cdk4*^*-/-*^ MEFs show increased fatty acid oxidation capacity. Interestingly, in hepatocytes, CDK4 controls glucose production through the general control non-repressed protein 5 (GCN5)-mediated inactivation of the peroxisome proliferator-activated receptor gamma coactivator 1-alpha (PGC-1α) in response to insulin (Lee, Nature 2014). The transcriptional coactivator PGC-1a plays a central role in the metabolism of a number of tissues, including skeletal muscle, by binding to transcription factors such as peroxisome proliferator-activated receptors (PPARs), estrogen-related receptors (ERRs) and nuclear respiratory factors (NRFs), and subsequently activating expression of nuclear and mitochondrial genes (Puigserver *et al*, 1998) (Scarpulla, 2011). Importantly, AMPK and PGC-1a are of utmost importance in the control of skeletal muscle oxidative metabolism and fiber type specification, and their activation promotes exercise-like adaptations including increased endurance (Lin *et al*, 2002; Narkar *et al*, 2008). Noteworthy, AMPK can also activate PGC-1a directly by phosphorylation (Jager *et al*, 2007).

Collectively, these results reported in different cell types and tissues show that CDK4 is able to repress the activity of important regulators of oxidative energy metabolism, which were previously shown to be crucial for skeletal muscle function. As *cdk4*^*-/-*^ mouse phenotype is characterized by leanness and endurance, we hypothesized that CDK4 regulates skeletal muscle physiology and therefore has an impact on the metabolism of the whole organism. Indeed, *cdk4*^*-/-*^ mice have higher fatty acid oxidation during exercise. These whole-body adaptations rely on skeletal muscle remodeling characterized by an increased mitochondrial oxidative capacity and a glycolytic-to-oxidative fibre switch. Transcriptomic analysis of *cdk4*^*-/-*^ gastrocnemius muscle (GNM) reveals the activation of the ERRα transcription factor, which is a well-described partner of PGC1α. This observation was validated using in vitro cultured C2C12 myotubes treated with a CDK4 inhibitor. Consistent with our previous findings (Lopez-Mejia, Mol cell 2017), skeletal muscle AMPK activation is augmented in the absence of CDK4. As it was described that AMPK phosphorylates PGC1α (Jager *et al*., 2007)and that PGC1α co-activates ERRα ((Huss *et al*, 2002; Schreiber *et al*, 2003), we propose that CDK4 deletion promotes ERRα transcriptional activity through AMPK-mediated activation of PGC1α. Finally, mice lacking CDK4 display an increased basal metabolic rate resulting in resistance to body weight gain and fat accumulation. Taken together, our data uncover a new function of CDK4 in the regulation of the oxidative capacity of skeletal muscle, ultimately resulting in changes in whole-body metabolism and protecting from body fat accumulation.

## Results

### Endurance and fat oxidation during exercise are increased in cdk4^-/-^ mice

In agreement with our previous findings, *cdk4*^*-/-*^ mice performed better during a graded running test (Figure 1A). Moreover, the hanging time was strikingly increased in *cdk4*^*-/-*^ mice during a hanging test (Figure 1B) and the total work, calculated as the product of hang time and body mass, remained significantly higher in *cdk4*^*-/-*^ mice (Figure 1C). This demonstrated that the energy production capacity in skeletal muscle of mice lacking CDK4 is improved independently of differences in body mass and excluded the possibility that the improved running capacity was simply the result of a lower body weight.

**Figure 1:**
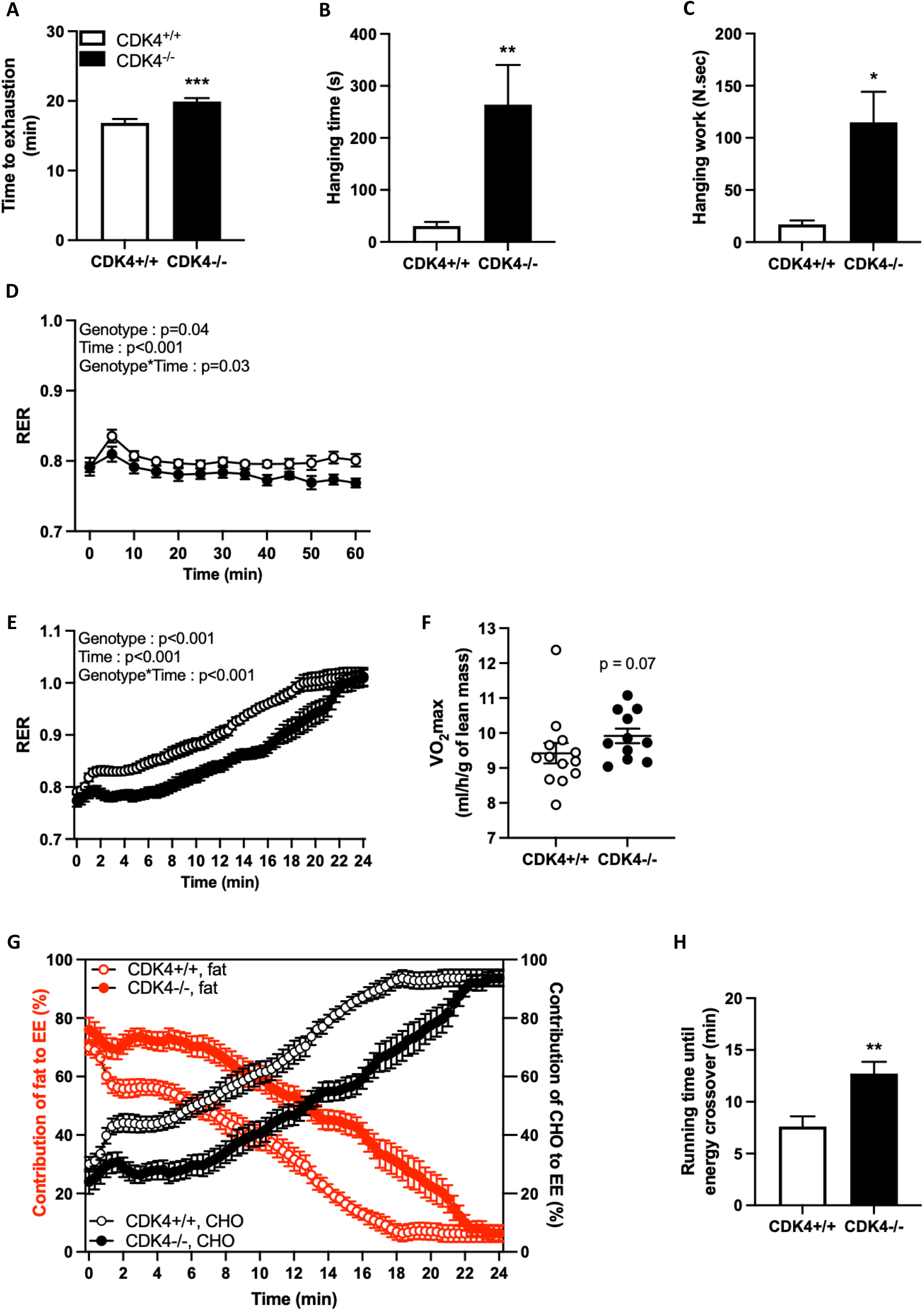
Endurance and fat oxidation during exercise are increased in *cdk4*^*-/-*^ mice. *Cdk4*^*+/+*^ and *cdk4*^*-/-*^ male mice were tested for exercise capacity between 20 and 25 weeks of age. (A) Time spent on the treadmill until exhaustion during a graded running test (n=11-13). (B-C) Time spent until fall and total work performed during hanging test (n=6-9). (D) Respiratory exchange ratio (RER) during a 1-hour, low and constant intensity running test (n=11-13). (E) Respiratory exchange ratio (RER) and (F) maximum oxygen uptake (VO_2_max) during a graded running test (n=11-13). (G) Relative contribution of fat (red) and carbohydrates (CHO, black) to total energy expenditure (EE) during a graded running test (n=11-13). (H) Running time until energy crossover measured from data presented in Figure 2G (n=11-13). Empty circles and bars: *cdk4*^*+/+*^; full circles and bars: *cdk4*^*-/-*^. *: p<0.05, **: p<0.01 and ***: p<0.001 for *cdk4*^*-/-*^ vs. *cdk4*^*+/+*^.

Next, to determine whether adaptations in energy metabolism in the *cdk4*^*-/-*^ mice could underline the higher endurance capacity, a low intensity (low and constant treadmill speed running for 1h) exercise was performed in combination with calorimetric measurements. No differences in blood lactate between rest and post-exercise conditions were observed during the low intensity run between genotypes (Figure Supp 1A), which confirmed that the exercise protocol is indeed moderate and does not force a metabolic switch to anaerobic glycolysis. Basal RER was also unchanged between *cdk4*^*+/+*^ and *cdk4*^*-/-*^ mice. In contrast, as exercise started, mice lacking CDK4 had a lower RER that became significantly decreased compared to *cdk4*^*+/+*^ mice after 40 minutes of run (Figure 1D). Accordingly, in *cdk4*^*-/-*^ mice the oxidation of carbohydrates and fat per unit of lean mass were decreased and increased, respectively, after 1h-run (Figure Supp 1C and 1D).

To test the ability of *cdk4*^*-/-*^ and *cdk4*^*+/+*^ mice to use fat at higher exercise intensities, which is a hallmark of training and is associated with higher endurance (Klein *et al*, 1994), we submitted these mice to a graded exercise (progressively increasing treadmill speed until exhaustion), coupled to a calorimetry test. Shortly after exhaustion, mice of both genotypes showed a similar burst of blood lactate values compared to the resting condition and the low-intensity exercise values (Figure Supp 1D) while most of the mice reached maximal RER values higher to 1 (Figure Supp 1E). Those results suggested that mice ran at their maximal capacity and highlighted the suitability of this protocol to push the animals to exhaustion. In line with the results obtained during the low-intensity exercise, the RER was consistently lower in *cdk4*^*-/-*^ mice until the expected final sharp increase up to 1 prior to exhaustion (Figure 1E). Consistent with the augmented endurance of *cdk4*^*-/-*^ animals, maximal oxygen consumption tended to be higher (Figure 1F). The relative contribution of fat and carbohydrates to total energy expenditure during the running test until exhaustion further showed that *cdk4*^*-/-*^ mice rely more on lipids as a source of energy during sub-maximal exercise (Figure 1G). The intersection of the carbohydrates (CHOs) and fat curves of each group of mice is named “energy crossover” and determines the time point when CHOs become the major (> 50%) source of energy. We illustrated in the figure 1H the differences in the time of the energy crossover between *cdk4*^*+/+*^ and *cdk4*^*-/-*^ mice, indicative of an increased capacity to preserve fatty oxidation as energetic sources in mice lacking CDK4 as exercise intensity increases.

Collectively, our results demonstrate that *cdk4*^*-/-*^ mouse endurance is augmented compared to their control littermates, relying on their improved capacity to oxidize fat during exercise. Since skeletal muscle is the major tissue contributing to changes in energy metabolism during exercise, these data suggest major adaptations in the skeletal muscle of *cdk4*^*-/-*^ mice, supporting higher oxidative capacities.

### Oxidative fiber type in the skeletal muscle of *cdk4*^*-/-*^ mice

We next focused on the morphological and molecular changes that could underlie the differences in endurance in *cdk4*^*-/-*^ mice. First, we isolated extensor digitorum longus (EDL) and soleus (SOL) muscles to measure maximal force and fatigability. EDL and soleus of *cdk4*^*-/-*^ mice had increased specific maximal forces (Figure Supp 2A and 2B), demonstrating an improvement in contractile function despite their smaller length and weight. These findings were consistent with the elevated force that *cdk4*^*-/-*^ mice developed during a grip strength test (Figure Supp 2C). The fatigability of the EDL muscle, which has a glycolytic metabolism, was similar in both genotypes (Figure Supp 2D and 2E), whereas the soleus, which has an oxidative metabolism, was more resistant to fatigue in muscles isolated from *cdk4*^*-/-*^ mice. Indeed, the *cdk4*^*-/-*^ soleus muscles showed a consistent higher percentage of their maximal force after repeated tetanic contractions (Figure 2A-2B). These results obtained in isolated muscles, *i*.*e*., with an experimental setting independent of any systemic acute regulation (namely nervous or circulating factors), support the hypothesis that increased endurance and resistance to fatigue result from the absence of CDK4 specifically in skeletal muscle.

**Figure 2:**
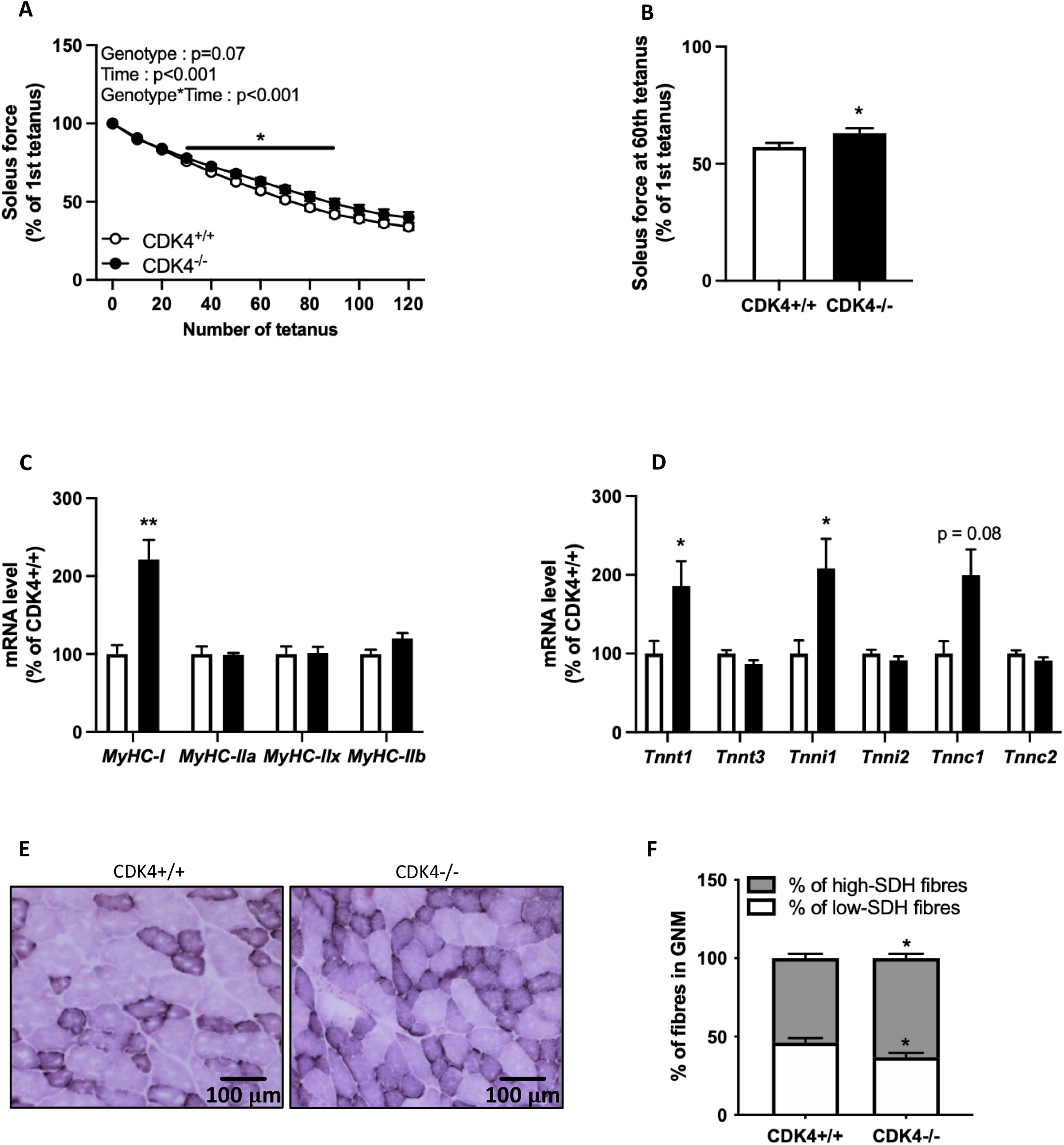
*Cdk4*^*-/-*^ mice display a skeletal muscle oxidative fibre type. *Cdk4*^*+/+*^ and *cdk4*^*-/-*^ male mice were euthanised at the age of 25 weeks. (A-B) Soleus muscle fatigability was assessed by measuring maximal force production along the course of repeated tetanic contractions (n=14-16). (C-D) mRNA levels of genes coding for slow and fast isoforms of myosin heavy chains (C) and troponin subunits (D) were measured by qPCR (n=5-7). (E) Transversal slices of GNM muscles were used to measure succinate dehydrogenase (SDH) activity, a marker of mitochondrial oxidative capacity. (F) Proportion of GNM muscle fibers stained negatively (SDH-neg) or positively (SDH-pos) for SDH activity (n=5). Empty circles and bars: *cdk4*^*+/+*^; full circles and bars: *cdk4*^*-/-*^. *: p<0.05 and **: p<0.01 for *cdk4*^*-/-*^ vs. *cdk4*^*+/+*^.

We further analyzed the muscle physiology by determining the fiber typology quantifying the expression of genes encoding for contractile proteins such as myosin heavy chain and troponin. The mRNA levels of *Myh7*, which is typical of slow, oxidative muscle fibers, was upregulated in the GNM of *cdk4*^*-/-*^ mice, while the expression of other myosin heavy chain isoforms characteristic of fast-twitch fibers was not changed (Figure 2C). Similarly, slow isoforms of troponin T, I and C (*Tnnt1, Tnni1* and *Tnnc1*) were upregulated in GNM lacking CDK4 while fast isoforms C (*Tnnt3, Tnni2* and *Tnnc2*) were not differentially expressed (Figure 2D).

Consistent with these functional and molecular features evoking a more oxidative phenotype, succinate dehydrogenase (SDH) activity, which is a marker of muscle fiber mitochondrial oxidative activity, was also increased in *cdk4*^*-/-*^ mice (Figure 2E and 2F). The experiments in TA muscles showed similar results and underscored that the oxidative phenotype associated with CDK4 deletion is not specific to a particular muscle (Figure Supp 2F and 2G).

Taken together, these results showed that CDK4 deletion promotes a muscle fiber switch towards a type 1, oxidative phenotype, sustaining the observed higher endurance in *cdk4*^*-/-*^ mice. Such functional and morphological changes require concomitant cellular metabolic adaptations, which prompted us to investigate the mitochondrial morphology and function.

### Cdk4^-/-^ mice have increased muscle mitochondrial content and oxidative capacity

The protein level of the outer mitochondrial membrane protein, TOMM20, was markedly elevated in mice lacking CDK4 (Figure Supp 3C), suggesting higher mitochondrial content in *cdk4*^*-/-*^ mice. Electron microscopy images of GNM longitudinal sections showed a higher percentage of cell volume occupied by mitochondria in the muscles of *cdk4*^*-/-*^ mice (Figures 3A and 2B). The increased mitochondrial volume was the result of a slight increase in the number (Figures 3A and 3C) and a marked augmentation of the size of mitochondria (Figures 3A and 3D). In addition to bigger size, the muscle mitochondria of *cdk4*^*-/-*^ mice also displayed changes in their shape. The aspect ratio, which is the quotient between the major and minor diameters, was higher in the absence of CDK4 (Figures 3A and 3E). Similarly, the form factor, which is a combined measure of mitochondrial length and degree of branching was also increased in *cdk4*^*-/-*^ muscles (Figures 3A and 3F). These morphological changes are indicative of mitochondrial elongation and interconnectivity (Harmuth *et al*, 2018; Koopman *et al*, 2005) and suggest a switch of mitochondrial dynamics towards a more fused state. Indeed, the activating phosphorylation of DRP1 (Ser617) (a major regulator of mitochondrial fission) was decreased, while the DRP1 inhibiting phosphorylation (Ser637) was increased (Figures 3G and 3H) in protein extracts from muscles of *cdk4*^*-/-*^ mice, further supporting a fused state consecutive to the inhibition of fission in these mice.

**Figure 3:**
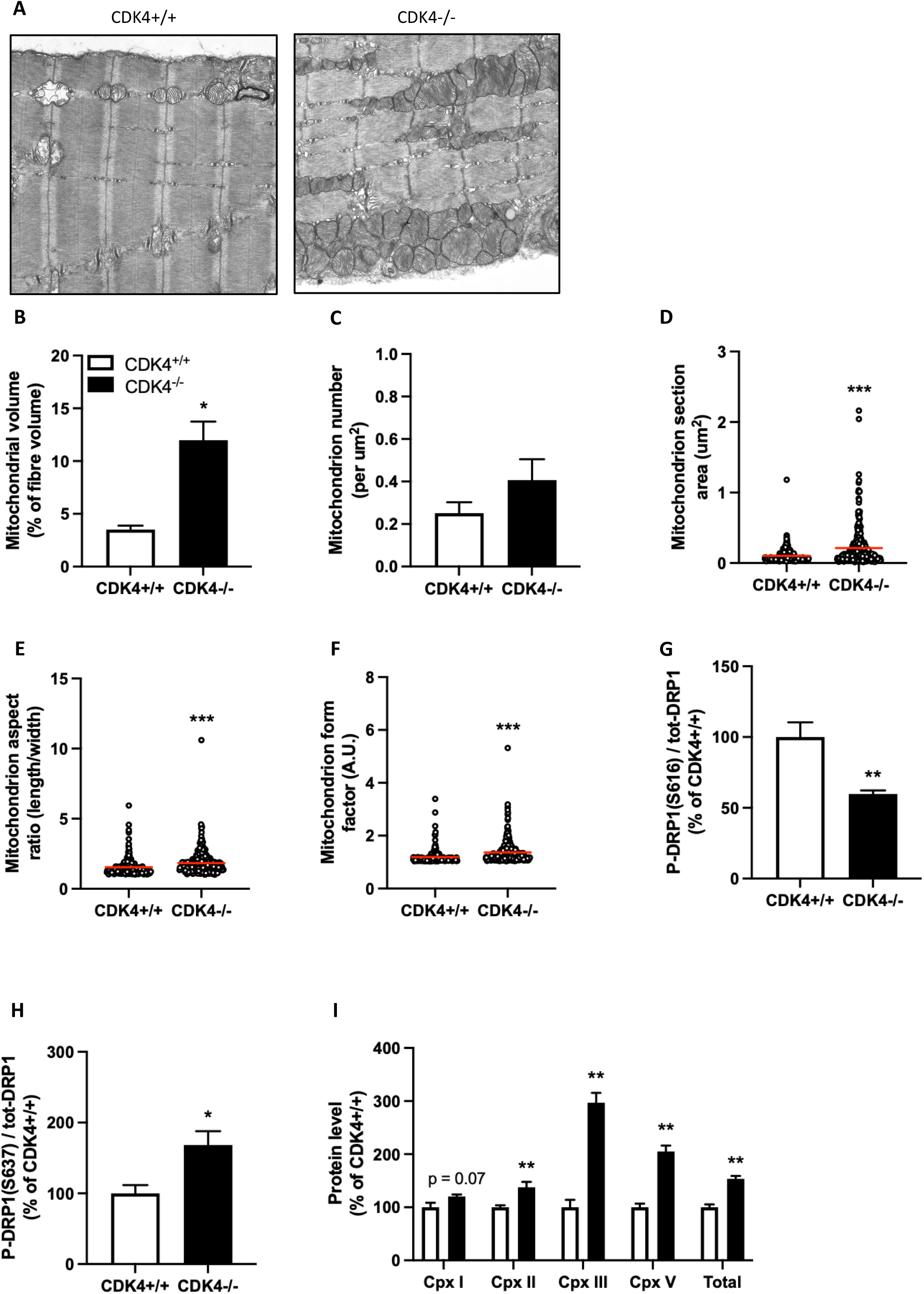
*Cdk4*^*-/-*^ mice have increased muscle mitochondrial content and oxidative capacity. *Cdk4*^*+/+*^ and *cdk4*^*-/-*^ male mice were euthanised at the age of 25 weeks. (A) Longitudinal slices of GNM were analysed by transmission electron microscopy. (B-F) Mitochondrial volume (B), number (C), section area (D), aspect ratio (E, index of elongation) and form factor (F, index of banching and complexity) (n=4). (G-H) Protein levels of P-DRP1 on Ser616 (G, inhibitory phosphorylation) and Ser637 (H, activating phosphorylation) normalised to total-DRP1, measured by western blot (n=6). (I) Protein levels of mitochondrial electron transport chain complexes measured by western blot. Empty circles and bars: *cdk4*^*+/+*^; full circles and bars: *cdk4*^*-/-*^. *: p<0.05, **: p<0.01 and ***: p<0.001 for *cdk4*^*-/-*^ vs. *cdk4*^*+/+*^.

The observed morphological changes in the mitochondria are evocative of post-exercise adaptations (Lundby & Jacobs, 2016; Tanaka *et al*, 2020) that come along with the upregulated expression of proteins of the respiratory chain complexes in *cdk4*^*-/-*^ mice (Figure 3I), further supporting an increase in mitochondrial oxidative capacity. Importantly, the increased oxidative capacity was not followed by oxidative damages (Figure Supp 3D), suggesting that these mitochondria are functional.

These results demonstrate that mice lacking CDK4 develop a mitochondrial phenotype evoking the training-induced adaptations, including morphological changes (increased mitochondrial content and elongation) and increased expression of metabolic genes, such as the subunits of the mitochondrial electron transport chain.

### Cdk4 ^-/-^ skeletal muscles display an ERR-dependent transcriptomic signature

To unravel the molecular changes underlining the improved endurance and skeletal muscle oxidative capacity in *cdk4*^*-/-*^ mice, we analyzed the gene expression profile of the GNM in resting conditions by RNAseq. More than 40 KEGG pathways were found to be significantly enriched among genes that were upregulated in the muscle of *cdk4*^*-/-*^, compared to *cdk4*^*+/+*^ mice (Figure 4A). Interestingly, some of the top-ranking pathways referred to energy metabolism, including both glucose- and fatty acid-related pathways (Figure 4A). Noteworthy, “oxidative phosphorylation” appeared as one the most significantly enriched pathways, further supporting the in vivo and the histological findings showing that *cdk4*^*-/-*^ mice have higher oxygen consumption and higher mitochondrial content in skeletal muscle.

**Figure 4:**
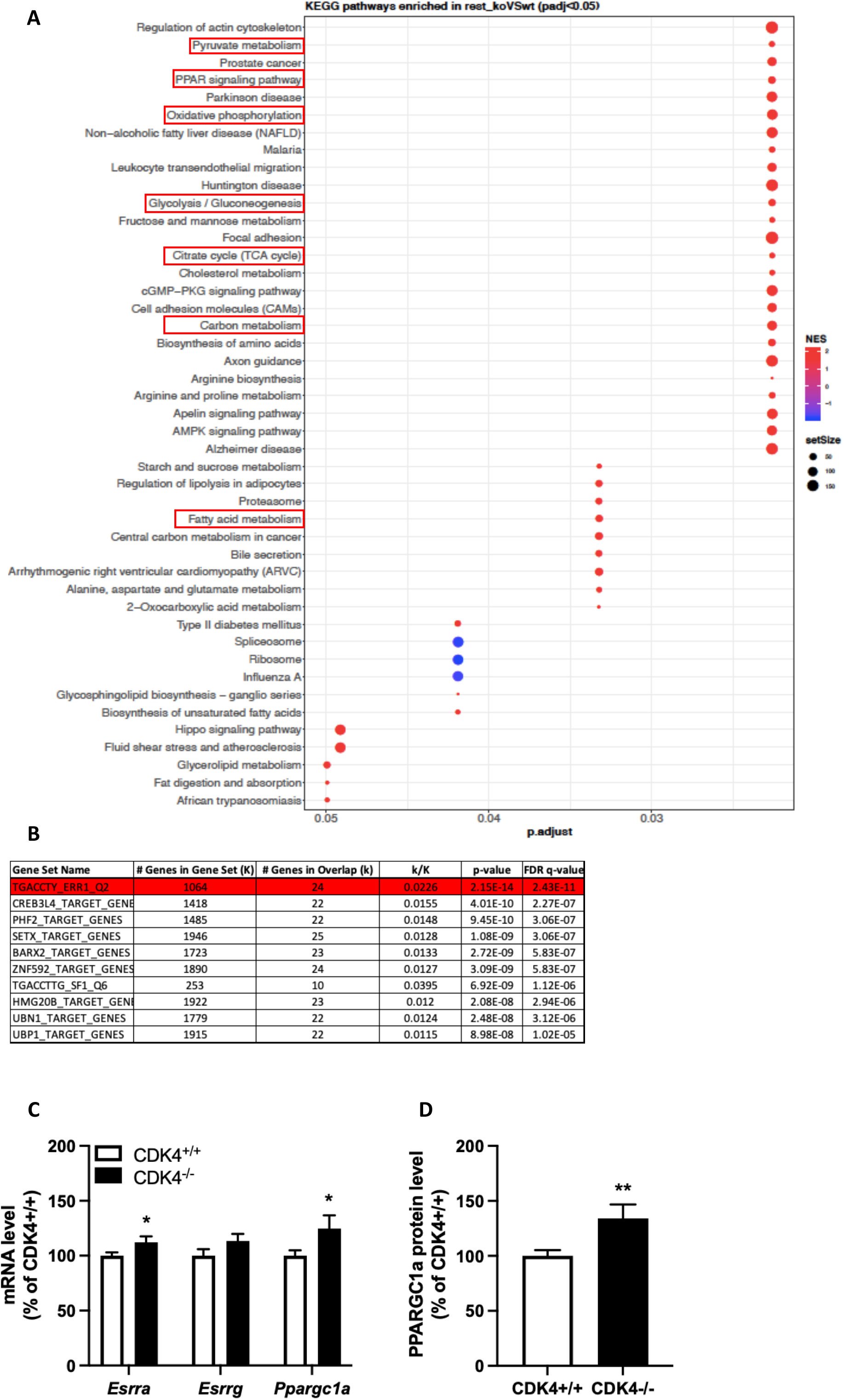
*Cdk4*^*-/-*^ skeletal muscles display an ERR-dependent transcriptomic signature. *Cdk4*^*+/+*^ and *cdk4*^*-/-*^ male mice were euthanised at the age of 30 weeks. (A) KEGG pathways significantly enriched in CDK4-/- vs CDK4+/+ GNM after the GSEA analysis was run against the KEGG pathway collection. Top pathways have the lowest adjusted p values (n=5-6). NES: normalised enrichment score. (B) Genes belonging to the top-ranking KEGG pathways related to energy metabolism (red squares on Figure 5A) were uploaded in the MsigDB collection to identify sets of genes including response elements to one single transcription factor. Top-ten gene set names and their associated transcription factors with the lowest adjusted q value are represented. (C-D) GNM mRNA (C) and protein (D) levels of the transcription factor ESRRa and its co-activator PPARGC1a (n=9-13). Empty bars: *cdk4*^*+/+*^; full bars: *cdk4*^*-/-*^. *: p<0.05 and **: p<0.01 for *cdk4*^*-/-*^ vs. *cdk4*^*+/+*^.

Seeking to decipher the molecular mechanism causing these transcriptional changes, we pooled the genes related to energy metabolism that were enriched in KEGG pathway analysis (Figure 4A, red boxes). This list was uploaded in the MSigDB collection “regulatory target gene sets-all transcription factor targets” to identify transcription factors likely to bind common response elements found in the promoters of these metabolic genes. The top-ranking cluster of genes contained a binding site for estrogen-related receptor alpha (*Esrrα* or *Errα*) in their regulatory region (Figure 5B), suggesting that ERRα activity is increased in skeletal muscle of *cdk4*^*-/-*^ mice. Strikingly, the gene expression of ERRα and PGC1α, which is a transcriptional target and coactivator of ERRα, was also upregulated in these mice (Figure 4C and 4D), while mRNA levels of *Errγ*, which shares most of its target genes with ERRα and is similarly co-activated by PGC1α, were slightly but not significantly increased (Figure 4C). qPCR analyses also showed an overall increase in the expression of genes coding for subunits of the mitochondrial respiratory chain (Figure Supp 4A).

**Figure 5:**
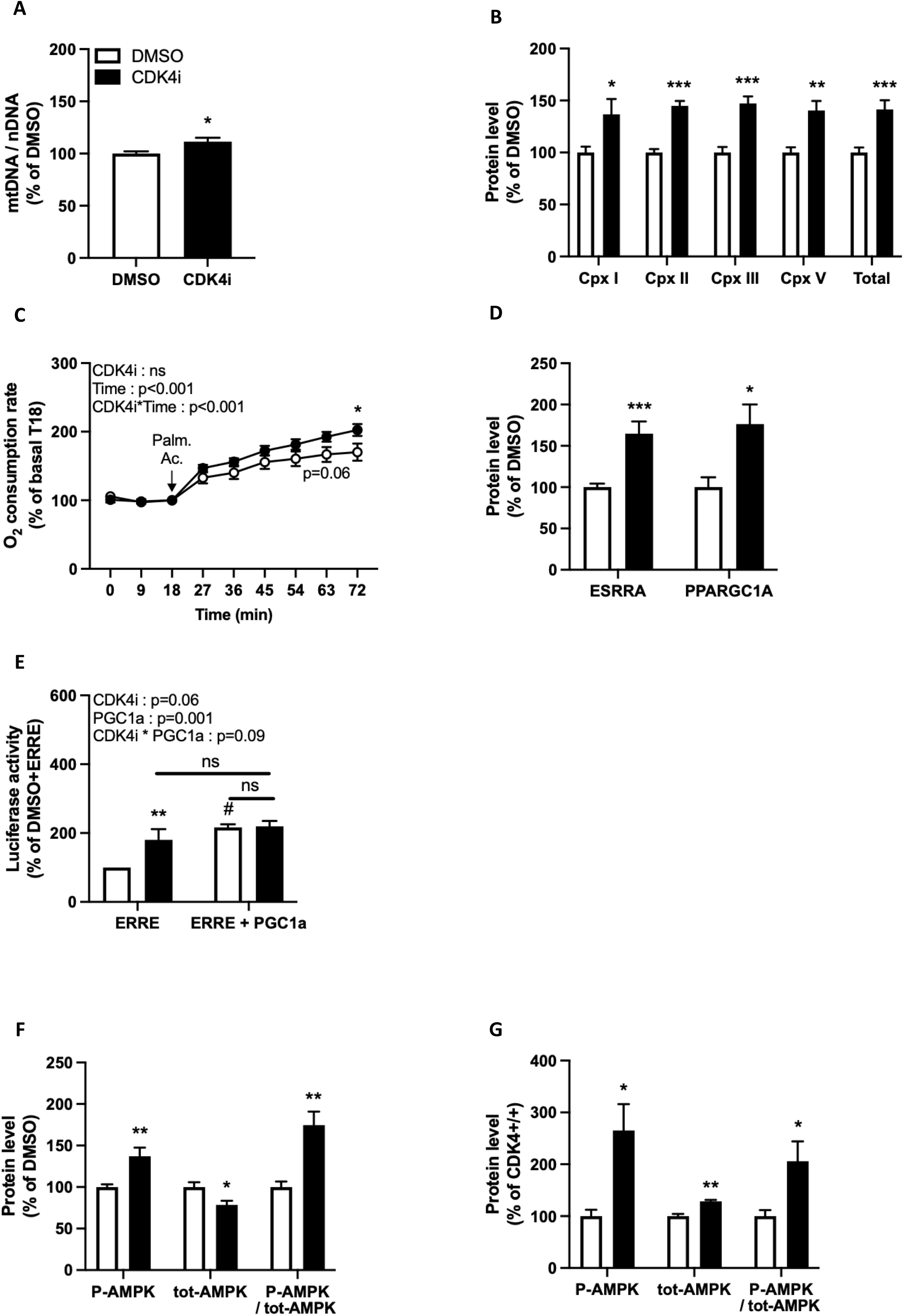
CDK4 inhibition in C2C12 myotubes stimulates ERR transcriptional activity. C2C12 myotubes were treated for 48h with abemaciclib, a CDK4/6 inhibitor (CDK4i). (A) Mitochondrial DNA (mtDNA) content normalised to nuclear DNA (nDNA) (n=9). (B) Protein levels of mitochondrial electron transport chain complexes measured by western blot (n=8). (C) Oxygen consumption rate measured by Seahorse in basal conditions (0-18 min) and after injection of palmitic acid (27-72 min) (n=8-10). (D) Protein levels of ERRa and PGC1α measured by western blot (n=8). (E) Luciferase activity measured by luminescence in C2C12 myotubes transfected with either only the ERR-reporter plasmid or with the ERR-reporter and a PGC1α-overexpressing plasmids(n=5-8). (F-G) Protein levels of Thr172 P-AMPK, total AMPK (tot-AMPK) and the ratio P-AMPK / tot-AMPK measured by western blot in C2C12 myotubes treated with DMSO or CDK4i (F) (n=6) and in CDK4+/+ and CDK4-/- GNM (G)(n=5-7). Empty circles and bars: DMSO (CDK4+/+ in panel 6G); full circles and bars: CDK4i (CDK4-/- in panel 6G). *: p<0.05, **: p<0.01 and ***: p<0.001 for CDK4i vs. DMSO (CDK4-/- vs. CDK4+/+ in panel 6G). #: p<0.05 for DMSO/ERRE+PGC1α vs. DMSO/ERRE.

These results suggested that the metabolic reprogramming of *cdk4*^*-/-*^ mouse skeletal muscle is mediated by the induction of the PGC1a/ERR transcriptional activity, ultimately resulting in the increased mitochondrial oxidative capacity required for the improved endurance capacity observed in *cdk4*^*-/-*^ mice.

### CDK4 inhibition in C2C12 myotubes stimulates ERR transcriptional activity

To further elucidate the molecular changes induced by the inhibition of CDK4 in muscle cells, we treated C2C12 myotubes for 48h with the CDK4/6 inhibitor (CDK4i) abemaciclib. The efficiency of the treatment was validated by the reduction in RB phosphorylation (Figure Supp 6A), which is the canonical target of CDK4 and a surrogate marker of its activity. CDK4i-treated cells revealed a higher mitochondrial DNA content (Figure 5A) and increased content of proteins of the electron transport chain (Figure 5B). Consistently, fatty acid-induced oxygen consumption was significantly higher in CDK4i-treated cells (Figure 5C). Similar to what we found in the skeletal muscle, ERRα and PGC1α mRNA and protein levels were upregulated in CDK4i-treated myotubes (Figure 5D and Figure Supp 5B). Noteworthy, ERRγ expression, which was barely detectable in DMSO-treated cells, was strongly increased upon CDK4i treatment (Figure Supp 5B). Importantly, increased ERR expression was accompanied by an increase in the transcriptional activity of ERR, as assessed by a luciferase assay where luciferase expression is under the control of an ERR response element (Figure 5E). Remarkably, the overexpression of PGC1α stimulated the ERR activity in DMSO-treated C2C12 myotubes but failed to further increase ERR activity upon CDK4i treatment (Figure 5E). These results suggested that the upregulation of the expression of PGC1α that we observed in CDK4i-treated cells could be induced by, and contribute to the increased ERR transcriptional activity, through a positive autoregulatory loop between PGC1α and ERRs (Schreiber *et al*., 2003). Accordingly, when ERRα is knocked down (Figure Supp 5C), the CDK4i-induced expression of PGC1α and ERRγ was, at least in part, abolished (Figure Supp 5D and 5E).

To verify the specificity of the abemaciclib treatment, we knocked down CDK4 by siRNA transfection in C2C12 myotubes. The resulting changes in gene expression were similar to what we observed upon CDK4i treatment, supporting a specific role of CDK4 in this process (Figure Supp 5F).

These results suggested that CDK4 blocks the transcriptional activity of the PGC1α-ERRα complex. PGC1α function is regulated at the post-transcriptional level by AMPK phosphorylation, which increases the activity of this nuclear receptor-coactivator. Strikingly, in C2C12 myotubes treated with abemaciclib and in gastrocnemius muscle of *cdk4*^*-/-*^ mice, the phosphorylation of T172-AMPK was significantly increased (Figure 5F and 5G), supporting the role of AMPK to mediate the effects of CDK4 inactivation on the PGC1α/ERR-dependent metabolic program.

### Cdk4 ^-/-^ mice have decreased body weight and increased basal metabolic rate

Body fat accumulation occurs when energy intake exceeds energy expenditure, ultimately leading to obesity and obesity-associated disorders such as insulin resistance. Hence, exercise and exercise-mimicking strategies aimed at increasing skeletal muscle oxidative capacity and energy expenditure have an anti-obesity potential. Interestingly, body weight gain at adult age is very limited in *cdk4*^*-/-*^ mice, with a disproportional decrease in fat mass accumulation (Figure 6A). This default in adipose tissue fat storage did not result in the accumulation of lipids in other organs, as evidenced by a lower fat content in liver and skeletal muscle (Figure 6C and 6D). Therefore, *cdk4*^*-/-*^ mice are not lipodystrophic but may have an imbalance between energy intake and energy expenditure.

**Figure 6:**
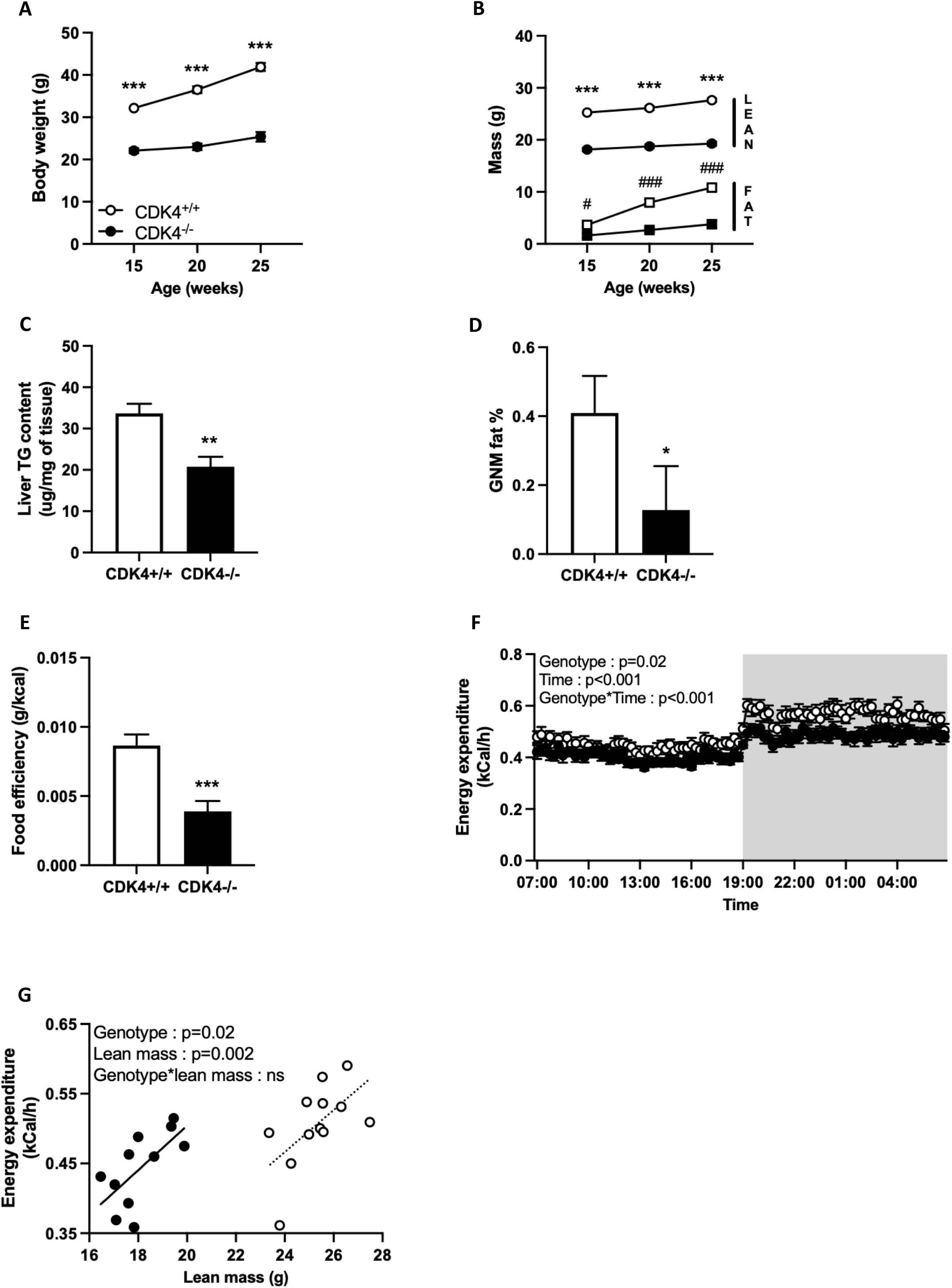
*Cdk4*^*-/-*^ mice have decreased body weight and increased metabolic rate. *Cdk4*^*+/+*^ and *cdk4*^*-/-*^ male mice were followed from 15 to 25 weeks of age. (A-B) Body weight and body composition (n=10-13). (C-D) Liver triglyceride content and gastrocnemius muscle fat content assessed by biochemical assay and echoMRI, respectively, in 25-week-old males (n=9-13). (E) Average food efficiency between 15 and 25 weeks of age (n=10-13). (F) Energy expenditure was calculated from indirect calorimetry experiments during the light (no background) and dark (grey background) phases over a 2-day period (n=11-12). (G) Average daily energy expenditure values were analyzed with the free online CalR application using lean mass as a covariable (n=11-12). Empty circles and bars: *cdk4*^*+/+*^; full circles and bars: *cdk4*^*-/-*^. ns: not significant, *: p<0.05, **: p<0.01 and ***: p<0.001 for *cdk4*^*-/-*^ vs. *cdk4*^*+/+*^.

Indeed, although body weight was strongly decreased in *cdk4*^*-/-*^, food intake was only slightly diminished (Figure Supp 6A). However, the relative food intake normalized to body weight was higher in *cdk4*^*-/-*^ mice (Figure Supp 6B), suggesting that decreased body weight and fat mass do not arise from defaults in feeding behavior. Decreased weight gain along with increased relative energy intake resulted in lower food efficiency (Figure 6E), which may originate from impaired energy absorption during digestion or increased energy expenditure in *cdk4*^*-/-*^ mice. No differences were found neither in the percentage of energy intake lost in the feces, nor in spontaneous locomotor activity between *cdk4*^*+/+*^ and *cdk4*^*-/-*^ mice (Figure Supp 6C and 6D). To further analyze energy homeostasis in these mice, we accurately evaluated the metabolic rate considering the differences in their body composition. Indeed, lean mass has to be taken into account as a covariable to analyze calorimetric data using the CalR software ((Mina *et al*, 2018; Speakman, 2013; Tschop *et al*, 2011). VO^2^ consumption, CO^2^ production and energy expenditure were significantly increased in *cdk4*^*-/-*^ compared to *cdk4*^*+/+*^ mice (Figures 6G and Supp 6F), demonstrating a higher basal metabolic rate in mice lacking CDK4. Interestingly, the respiratory exchange ratio (RER) was similar in both genotypes (Figure Supp 6G), indicating that the relative contribution of glucose and fatty acids as energetic substate is not different in the absence of any metabolic stress.

Collectively, our results show that *cdk4*^*-/-*^ mice are protected from fat accumulation by an increase in energy expenditure. Although every organ has a specific metabolic rate, the main body compartment contributing to total energy expenditure is lean mass, which is predominantly made of skeletal muscle. Together with our observations that *cdk4*^*-/-*^ mice are more endurant and show specific skeletal muscle adaptations, our results highlight muscle CDK4 as a potential target to counteract obesity development.

## Discussion

Changes in substrate utilization is a major tool of the organism to adapt to specific needs. For instance, during endurance training skeletal muscles shift their metabolism from glucose to fatty acids as a main energy source. In this work we show that the cell cycle regulatory protein CDK4 is a key kinase that controls the energy balance in muscle cells, resulting in an overall impact on the whole organism metabolism. Along with improved muscle performance, energy expenditure is increased in mice lacking CDK4, thereby contributing to the reduced weight and fat accumulation. We demonstrate that the deletion of CDK4 causes profound changes in the metabolism of skeletal muscle, including a better capacity to use lipids as energy substrates which is a feature of endurance training (Achten & Jeukendrup, 2004; Scharhag-Rosenberger *et al*, 2010). Consequently, *cdk4*^*-/-*^ mice show an increased endurance. Moreover, the improved contractile function underscores the tissue-specific resistance to fatigue of *cdk4*^*-/-*^ compared to *cdk4*^*+/+*^ isolated muscles. In addition, histological and molecular analyses of the skeletal muscle of *cdk4*^*-/-*^ mice, reveal a higher mitochondrial content and metabolic adaptation, reminding a fast-to-slow fiber type switch. Indeed, *cdk4*^*-/-*^ mice have a basal muscle phenotype that evokes molecular training effects, including upregulation of pathways and genes involved in fatty acid metabolism (Lundsgaard *et al*, 2018; Purdom *et al*, 2018) as well as higher mitochondrial content (Hood *et al*, 2011). Muscle-specific effects of CDK4 are further evidenced in vitro, as C2C12 myotubes treated with a CDK4 inhibitor showed an increase in fatty acid oxidation capacity, consistent with the calorimetry results showing that *cdk4*^*-/-*^ mice use preferentially fat as energy substrate during exercise. Lundby and Jacobs (Lundby & Jacobs, 2016)reported that mitochondrial content rises upon training as a consequence of an increase in the mitochondrial cross-sectional area and in longitudinal growth (i.e., elongation), which is similar to the mitochondrial morphology in CDK4-/- skeletal muscle. Elongated mitochondria are protected from autophagy, have more cristae and are more efficient in ATP production (Gomes *et al*, 2011), which provides an obvious metabolic advantage in the exercising muscle.

CDK4 deficiency recapitulates at some extent the function of the estrogen-related receptors (ERRα, and ERRγ) in skeletal muscle (Fan & Evans, 2015). ERRs are nuclear receptors which transcriptional activity is of utmost importance in the regulation of energy metabolism in skeletal muscle. Indeed, ERRα and ERRγ expression is enriched in slow, oxidative, compared to fast, glycolytic muscles (Huss *et al*, 2004) (Rangwala *et al*, 2010). ERR targets include fatty acid oxidation, tricarboxylic acid cycle and OXPHOS genes. We show here that the increase in the oxidative capacity of the muscles of *cdk4*^*-/-*^ mice is driven by the transcriptional activity of the nuclear receptors ERRs and their co-activator PGC1α (Huss *et al*., 2004) (Giguere, 2008). ERRγ may act synergistically with ERRα as both nuclear receptors have widely overlapping metabolic target genes and functions (Audet-Walsh & Giguere, 2015). Interestingly, PGC1α expression is upregulated in *cdk4*^*-/-*^ skeletal muscles and by CDK4i in muscle cells, and the ERR reporter activity is not further increased when PGC1α is co-expressed. These results suggest that PGC1α upregulation in the absence of CDK4 may drive the increase in ERR transcriptional activity. Inhibition of PGC1α activity by CDK4 was previously reported in hepatocytes (Lee *et al*, 2014), where activation of GCN5 triggered PGC1α acetylation to decrease the transcription of its target genes. A different mechanism is likely to be involved in skeletal muscle, where PGC1α upregulation and activation is a well-described consequence of exercise training (Goto *et al*, 2000; Wu *et al*, 2002) that is mediated, at least in part, by AMPK ((Jager *et al*., 2007; Zong *et al*, 2002)Interestingly, we previously reported that AMPK is a direct target of CDK4. CDK4 inhibits AMPK by phosphorylation of Ser377, associated with a default in the classical activating phosphorylation on Thr172 (Lopez-Mejia *et al*., 2017). In CDK4-/- skeletal muscle and C2C12 myotubes treated with CDK4i, Thr172 phosphorylation is higher, suggesting that AMPK mediates the beneficial effects of CDK4 inactivation in muscle through regulation of gene expression driven by PGC1α-dependent ERR transcriptional activity.

We cannot exclude, however, a complementary AMPK-independent effect of CDK4 on mitochondrial activity or dynamics. As a matter of fact, other members of the CDK family, such as CDK1 and CDK5 phosphorylate the dynamin-related protein 1 (DRP1) and activate mitochondrial fission (Taguchi *et al*, 2007) (Jahani-Asl *et al*, 2015). We can speculate that CDK4, in addition to regulating activity of the AMPK-PGC1α-ERR transcriptional pathway, phosphorylates DRP1 to activate mitochondrial fission. In support of this hypothesis, we show here that the activating phosphorylation of Drp1 is reduced in the *cdk4*^*-/-*^ muscle and, consequently, mitochondria are in an hyperfused state.

The kinase activity of CDK4 is regulated by the members of the cyclin D family. Interestingly, cyclin D3-knockout mice exhibit higher energy expenditure, lower RER, improved endurance, and a fiber type switch towards type I oxidative fibers in the skeletal muscle (Giannattasio *et al*, 2018), which is comparable to the phenotype of *cdk4*^-/-^ mice. This is fully consistent with the function of cyclin D3 in the activation of CDK4. Similarly, cyclin D1 was found to repress fatty acid oxidation in hepatocytes through inhibition of PPARα activity (Kamarajugadda *et al*, 2016), further highlighting the importance of cell cycle regulators in the control of oxidative metabolism.

The overall control of energy homeostasis is the result of the functions and interactions of distinct tissues including, but not limited to, the brain, adipose tissue, liver and skeletal muscle. The present study undoubtedly demonstrates that CDK4 deletion triggers a catabolic state evidenced by an increase in energy expenditure. We show here that in muscle, CDK4 depletion promotes mitochondrial activity through the activation of the AMPK-PGC1α-ERR pathway, which primes the cells for substrate oxidation, namely fatty acid during exercise. However, the sole skeletal muscle by itself cannot fully account for this phenotype, as energy substrates need to be supplied from other tissues to fuel its oxidative metabolism. Our finding that RER is not changed in *cdk4*^-/-^ mice in basal conditions indicates that the augmentation in energy expenditure is supported by both glucose and lipid consumption. Interestingly, we previously reported an increased lipolysis in explants of white adipose tissue, the main reserve of fat in the organism (Lagarrigue *et al*., 2016). On the other hand, Lee et al (Lee *et al*., 2014) showed that inactivation of CDK4 stimulates glucose production in liver, the main reserve of glycogen. Altogether, these results suggest that CDK4 may function to coordinate whole-body metabolism by inhibiting substrate delivery to, and energy consumption in, myofibres. In the absence of CDK4, systemic release of fatty acids and glucose sustain higher energy expenditure, resulting at the long-term in the healthy, lean phenotype of *cdk4*^-/-^ mice.

In normal-weight individuals, lean mass comprises more than 50% of muscle mass and is the main factor contributing to variations in daily energy expenditure (Muller *et al*, 2018). Nevertheless, the heterogeneity of metabolic rates of single tissues composing the lean mass (Muller *et al*., 2018; Speakman, 2013) does not allow determining whether changes in basal energy metabolism of *cdk4*^-/-^ mice are only mediated by skeletal muscle. Moreover, we have recently described that mice lacking CDK4 specifically in the ventromedial hypothalamus are more tolerant to cold. This was a consequence of increased thermogenic activity of brown adipose tissue and indicated a specific function of CDK4 in the brain to control peripheral metabolism, which may contribute to the increase in energy expenditure of *cdk4*^-/-^ mice (Castillo-Armengol *et al*, 2020). Interestingly, other laboratories also suggested that CDK4 plays an important role in the hypothalamus. Intracerebroventricular injection of the CDK4/6 inhibitor abemaciclib in diet-induced obese mice reduced fat mass and increased lipid oxidation (Iqbal *et al*, 2018). In another study, it was shown that the treatment of mice fed a high fat diet with abemaciclib also reduced the fat mass and body weight in these animals in a MC4R-dependent manner, highlighting the central effects of the inhibitor (Iqbal *et al*, 2021). The latter studies should be taken with caution, since abemaciclib is not a specific inhibitor of CDK4, but is also an inhibitor of CDK6, and to a lesser extent also of CDK9 (Hafner *et al*, 2019).

Taken as a whole, the function of CDK4 in the cell is to facilitate the biosynthetic processes that are required for the cell to divide in anabolic conditions, and therefore contribute to the growth of the cellular population. Translated to the whole organism, we can speculate that CDK4 has a similar role. Indeed, by blocking oxidative metabolism in muscle, as well as in other tissues such as brown adipose tissue (Castillo-Armengol *et al*., 2020), CDK4 preserves the energy stores from being used by exercise in the muscle and keep them for biosynthetic processes when needed. Therefore, repurposing of CDK4 inhibitors may be considered as a valuable therapeutic option to promote energy expenditure through increased muscle oxidative capacity, thereby preventing or alleviating obesity-induced metabolic disorders, including insulin resistance, or muscle dystrophies characterized by a decrease in mitochondrial function such as DMD (Handschin *et al*, 2007; Selsby *et al*, 2012; Timpani *et al*, 2015). Importantly, *cdk4*^-/-^ mice fat content is diminished not only in adipose tissue, but also in liver and skeletal muscle where ectopic lipid deposition participates in the development of metabolic diseases such as non-alcoholic fatty liver disease or type 2 diabetes (Bosma *et al*, 2012; Loomba *et al*, 2021).

In conclusion, our study uncovers the role of CDK4 in the control of whole-body energy metabolism and exercise capacity through activation of the PGC1a-ERR axis in skeletal muscle. These findings suggest that CDK4 inhibitors could be considered as an exercise mimetic and may be a promising approach to prevent or treat obesity-associated diseases. As CDK4 deletion promotes oxidative metabolism in skeletal muscle, CDK4 inhibitors or gene therapy-mediated CDK4 knockdown may also be considered in the treatment of myopathies associated with impaired mitochondrial function such as Duchenne muscular dystrophy.

## Acknowledgements

The authors acknowledge all the members of the Fajas laboratory for support and discussions. The authors thank Sandra Calderon, Johann Weber and Hannes Richter from Lausanne Genomic Technologies Facility for RNA sequencing sample preparation and data analysis. We also thank Gilles Willemin from Mouse Metabolic Facility of CIG (University of Lausanne, Switzerland) for experimental assistance, and the Phenogenomics platform of EPFL (Lausanne). This work was supported by the Swiss National Foundation (179271). The authors thank Etienne Mouisel and Nathalie Didier for helpful discussion.

## Materials and methods

### Animals

Generation of CDK4+/+ and CDK4-/- mice were previously described (Martin, Oncogene 2003; CDK4-/- were referred to as CDK4^n/n;+/Cre^ in this original paper). Animals were constantly maintained in a temperature-controlled animal facility with a 12 h light/12 h dark cycle and free access to standard chow diet (SAFE-150, SAFE SAS) and water. Male mice were studied between the age of 15 and 25 weeks. All animal care and treatment procedures were performed in accordance with the Swiss Animal Protection Ordinance (OPAn) and were approved by the Canton of Vaud veterinary service (authorization VD3398h).

Body composition (fat and lean mass) was measured using EchoMRI technology with the EchoMRI^™^ 3-in-1 device (EchoMRI LLC) under isoflurane anesthesia. Indirect calorimetry and locomotor activity measurements were performed with an Oxymax/CLAMS system (Columbus Instruments, Columbus, OH, USA) with free access to food and water. The free online CalR program (Mina *et al*., 2018) was used to implement a generalized linear model (GLM) analysis utilizing lean mass as a covariate when modeling lean mass-dependent metabolic parameters using a two-group template. The results of this analysis are depicted in Figure 1H, where the significance test for “Genotype” is whether CDK4+/+ and CDK4-/- mice are significantly different for the metabolic variable selected, and the significance test for “Lean mass” determines whether there is an association between the lean mass and metabolic variables selected among all animals in the study. Feces energy content was measured by calorimetry bomb over a period of 3 days.

Running experiments were performed with a Treadmill Simplex II (Columbus Instruments) coupled to the above-mentioned indirect calorimetry system. Mice were acclimated to the treadmill on D-2 (10 min at 0 m/min, then 10 min at 10 m/min) and D-1 (10 min at 0 m/min, then 10 min at 12 m/min) prior to the tests. The day of the tests, mice were placed on the treadmill at 0 m/min for 15 min in order to stabilize gas exchanges and to decrease stress due to manipulation. During the low intensity running test, the speed of the treadmill was set at 8.5 m/min for 10 min, then 10 min at 10 m/min and 40 min at 12.5 m/min. During the graded test, mice started to run at 7 m/min for 1 min, then speed was increased by 0.5 m/min every 20 sec until exhaustion (Ayachi *et al*, 2016). Carbohydrate (CHO) and fat oxidation were calculated as follows (Jeukendrup & Wallis, 2005):

-low-intensity exercise CHO oxidation (g/min): 4.344 x VCO_2_ (l/min) – 3.061 x VO_2_ (l/min)

-graded exercise CHO oxidation (g/min): 4.210 x VCO_2_ (l/min) – 2.962 x VO_2_ (l/min)

-low-intensity and graded exercise fat oxidation: 1.695 x VO_2_ (l/min) – 1.701 x VCO_2_ (l/min)

Blood lactate was measured at rest and immediately after exercise from tail vein using a strip-based lactate analyzer (EDGE). VO_2_max was determined as the peak O_2_ consumption during graded exercise (as skeletal muscle is the major tissue contributing to O_2_ consumption during exercise, values of VO_2_ were normalized to lean mass), and maximal RER as the highest RER value reached during graded exercise including running time and 2-minute post-exhaustion. Energy crossover was determined as the time point at which contribution of CHO to energy expenditure exceeds fat contribution during a graded exercise.

The hanging test was performed by placing mice on a grid that was immediately switched upside-down. Each animal was submitted to 3 trials with a 5-minute in between periods of rest. Maximal time to fall was reported and holding impulse values were obtained by multiplying hang time (in seconds) by body weight (in Newtons).

A grip strength test device (Bioseb) made of a grid connected to a sensor was used to measure grip strength. Mice were held by the tail, allowed to grasp the grid with 4 paws then pulled back gently and steadily until the grid was released. The maximum force (in Newton) achieved by the animal before releasing the grid was recorded by the sensor. Each animal was given 3 trials with a 5-minute rest in between. The highest value was reported and normalized to body weight.

All in vivo experiments were implemented at the Mouse Metabolic Evaluation Facility (MEF) of the University of Lausanne.

### Isolated skeletal muscle contractile properties

Muscle mechanical measurements were assessed as previously described (Luan *et al*, 2021; Zanou *et al*, 2010) with slight modifications. All assays were measured in a blinded fashion. Twenty-five to 28-week-old CDK4+/+ and CDK4-/- mice were euthanized by cervical dislocation. Extensor digitorum longus (EDL) and soleus muscles were quickly dissected, then bathed in a 10ml horizontal chamber containing a continuously oxygenated Krebs solution composed of 135.5 mM NaCl, 5.9 mM KCl, 1 mM MgCl_2_, 2.5 mM CaCl_2_, 11.6 mM HEPES sodium, and 11.5 mM glucose, pH 7.4 at 25°C. The muscle was tied between a dual-mode lever arm and a fixed hook, and a stimulation was delivered through platinum electrodes running parallel to the muscle (1500A Intact Muscle Test System, Aurora Scientific Inc., Canada). Resting muscle length (*L*_*0*_) was carefully adjusted for maximal isometric force with 125-Hz maximally fused tetani. The force–frequency relationship was determined by sequentially stimulating the muscles at 25, 50, 75, 100, 125 and 150 Hz stimulation trains of 300ms duration with 1 min rest between each contraction. Normalized muscle specific force (mN/mm ^2^) was expressed relative to the cross-sectional area (CSA), obtained by dividing muscle blotted weight (mg) by muscle length and considering the fiber length equal to 0.5 L_0_ for EDL and 1 for soleus muscles (Brooks & Faulkner, 1988). To investigate muscle fatigue, muscles were subjected to 125 Hz stimulation trains of 300 ms duration at 10 s intervals over 50 tetanus for EDL muscles and at 1 s interval over 120 tetanus for soleus muscles. Data from each experiment were analyzed with Aurora’s DMA software (Aurora Scientific Inc., 2002, Solwood Enterprises, Inc., 2002) and Microsoft Excel.

### Skeletal muscle histology and immunofluorescence

GNM and TA were dissected from 25 week-old mice, embedded in OCT compound (Tissue-Tek) and frozen in liquid nitrogen-cooled isopentane. Mitochondrial oxidative capacity was assessed by measuring succinate dehydrogenase activity directly on muscle cryosections, through the intensity of nitro blue tetrazolium purple staining following succinate (the SDH substrate) addition. The percentage of single fibres categorized as “SDH-negative” or “SDH-positive” according to staining intensity were quantified on two fields of 5 mice per group (200-600 fibres per animal).

### Liver and skeletal muscle lipid content

A piece of snap-frozen liver was used for lipid extraction by a modified Folch method. Triglycerides were then quantified using a Cobas C111 analyser (Roche diagnostics), and results were normalized to sample weight.

Fat content was assessed in freshly dissected GNM by using EchoMRI technology with the “biopsy sample” option of the EchoMRI^™^3-in-1 device (EchoMRI LLC).

### Skeletal muscle transmission electron microscopy

After GNM muscles were dissected, pieces of 1mm^3^ were cut from the same region of the muscle belly and fixed in glutaraldehyde solution (EMS) 2.5% in Phosphate Buffer (PB 0.1M pH7.4) (Sigma-Aldrich) during 2h at room temperature (RT). Then, they were rinsed 3 times 5 minutes in PB buffer and then postfixed by a fresh mixture of osmium tetroxide 1% (EMS) with 1.5% of potassium ferrocyanide (Sigma-Aldrich) in PB buffer during 2h at RT. The samples were then washed three times in distilled water and dehydrated in acetone solution (Sigma-Aldrich) at graded concentrations (30%-40min; 70%-40min; 100%-1h; 100%-2h). This was followed by infiltration in Epon resin (Sigma-Aldrich) at graded concentrations (Epon 1/3 acetone-2h; Epon 3/1 acetone-2h, Epon 1/1-4h; Epon 1/1-12h) and finally polymerized for 48h at 60°C in the oven. Ultrathin sections of 50nm were cut on a Leica Ultracut (Leica Mikrosysteme GmbH) and picked up on copper slot grid 2×1mm (EMS) coated with a polystyrene film (Sigma-Aldrich). Sections were poststained with uranyl acetate (Sigma-Aldrich) 2% in H_2_O for 10 minutes, rinsed several times with H_2_O followed by Reynolds lead citrate for 10 minutes and rinsed several times with H_2_O. Micrographs were taken with a transmission electron microscope Philips CM100 (Thermo Fisher Scientific) at an acceleration voltage of 80kV with a TVIPS TemCam-F416 digital camera (TVIPS GmbH).

Micrographs with a pixel size of 4.1nm over an area of 100×100µm were taken with a transmission electron microscope Philips CM100 (Thermo Fisher Scientific) at an acceleration voltage of 80kV with a TVIPS TemCam-F416 digital camera (TVIPS GmbH) and the alignment was performed using Blendmont command-line program from the IMOD software (Kremer *et al*, 1996). Those large images were firstly sampled by systematic uniform random sampling using the Fiji plugin “unbiased counting frame”; then, in the identified areas (representing a total of 200 um^2^ per animal), mitochondrion number, volume and morphology were measured with Fiji (version 2.3.0).

### Skeletal muscle RNA sequencing

GNM muscles of mice euthanized in the morning in fed state were ground in liquid nitrogen and total RNA was extracted from the powder obtained using TRI Reagent (#T9424, Sigma-Aldrich), following manufacturer’s instructions. RNA quality was controlled by a Fragment Analyzer system (Agilent) and 6 samples per group with an RNA Integrity Number > 8 were used. RNA-seq libraries were prepared from 500 ng of total RNA with the Illumina TruSeq Stranded mRNA reagents (Illumina) using a unique dual indexing strategy, and following the official protocol automated on a Sciclone liquid handling robot (PerkinElmer). Libraries were quantified by a fluorimetric method (QubIT, Life Technologies) and their quality assessed on a Fragment Analyzer. Cluster generation was performed with 2 nM of an equimolar pool from the resulting libraries using the Illumina HiSeq 3000/4000 SR Cluster Kit reagents and sequenced on the Illumina HiSeq 4000 using HiSeq 3000/4000 SBS Kit reagents for 150 cycles (single end). Sequencing data were demultiplexed using the bcl2fastq2 Conversion Software (version 2.20, Illumina).

After filtration of lowly expressed genes and normalisation to the library size, a principal component analysis identified one CDK4-/- sample as an outlier that was further removed from the analysis. Thus, 6 CDK4+/+ and 5 CDK4-/- GNM samples were used for a Gene Set Enrichment Analysis (GSEA), in order to uncover overrepresented pathways within the entire list of expressed genes (>13,000) ranked according to the p value of their differential expression in CDK4-/- *vs*. CDK4+/+ animals. The GSEA results were then run against the KEGG pathway database, ignoring gene sets containing less than 10 genes or more than 500 genes.

### C2C12 cell culture

C2C12 myoblasts obtained from the American Type Culture Collection (ATCC) were grown in DMEM (#41966-052, Gibco) containing non-essential amino acids (#11140-076, Gibco) and 10% fetal bovine serum (FBS, #10082-147, HyClone). At confluence, FBS was replaced by 2% horse serum (#SH30074.03, HyClone). After 4 days of differentiation, cells were treated with 1.5uM of the CDK4 inhibitor (CDK4i) abemaciclib (LY2835219, MedChem Express) for 48 hours.

### Seahorse

C2C12 oxygen consumption rates (OCR) were analysed using an XFe96 Seahorse Analyzer (Seahorse bioscience). Cells were cultured as described above until the day of experiment. Differentiation medium was then replaced by a KHB medium containing 10mM glucose and 1.5mM carnitine and cells were incubated for 1 hour at 37°C without CO_2_ to allow for equilibration of the assay medium before starting the measurements. After measuring baseline OCR indicative of basal respiration, 300uM of palmitic acid coupled to BSA were injected to assess fatty acid oxidation. Data were normalised by protein content using a Pierce BCA protein assay kit (#23225, Thermo Fisher Scientific).

### Luciferase reporter

C2C12 myotubes were used at D4 of differentiation. Using Lipofectamine 3000 (Invitrogen), cells were transfected with 300 ug of a pGL2 plasmid expressing firefly luciferase under the control of a minimal promoter including 3 estrogen-related receptor response elements (ERRE); cells were simultaneously transfected with 50 ug of a pCMV5 plasmid expressing beta-galactosidase. One hundred and fifty micrograms of a pCDNA3 plasmid expressing PGC1α were also co-transfected in the corresponding wells. After overnight transfection, medium was replaced by differentiation medium. Fourty-eight hours post-transfection, cells were harvested in a Renilla luciferase assay buffer (#E290A, Promega) and lysed for 30 minutes at room temperature. Cell lysate was then assessed for luciferase and beta-galactosidase activities by luminescence and spectrophotometry, respectively. To control for transection efficiency, the luciferase signal was normalised by the beta-galactosidase activity in each well.

### siRNA transfection

C2C12 myotubes were used at D4 of differentiation. Using Lipofectamine RNAimax (Invitrogen), cells were transfected with 20nM of a mix of 2 control siRNAs (#SIC001 and #SIC002, Sigma-Aldrich), or 20nm of a mix of 2 siRNAs targeting Esrra (siEsrra-1, sense: AGGUGGUGGGCAUCGAGCCUCUCUA, antisense: UAGAGAGGCUCGAUGCCCACCACCU; siEsrra-2 sense: UGUCAGUACUGCAGAGUGU, antisense: ACACUCUGCAGUACUGACA; Sigma-Aldrich) or 20nm of a mix of 3 siRNAs targeting Cdk4 (siCdk4-1, sense: GGGAAAAUCUUUGAUCUCA, antisense: UGAGAUCAAAGAUUUUCCC; siCdk4-2 sense: CAGCACAGUUCGUGAGGUG, antisense: CACCUCACGAACUGUGCUG; siCdk4-3 sense: GCGCAGCUGCUACUGGAAA, antisense: UUUCCAGUAGCAGCUGCGC; Sigma-Aldrich). After overnight transfection, medium was replaced by differentiation medium and cells were harvested 48h post-transfection.

### qPCR on skeletal muscle and C2C12 myotube samples

RNA extraction from skeletal muscle was performed as described above (“Skeletal muscle RNA sequencing). For C2C12 myotubes, cells were harvested directly into TRI reagent and frozen prior to RNA extraction.

RNA concentrations were determined with a NanoDrop 8000 (Thermo Fisher Scientific) and reverse transcribed using 1,000 ng of RNA and Superscript II enzyme (18064014, Invitrogen) according to the manufacturer’s instructions. qPCR analysis was performed using SYBR Green detection (04913914001, Roche) on a QuantStudio 6 system (Applied Biosystems) according to the manufacturer’s instructions. Relative mRNA expression was calculated by the 2^^ (Delta Ct)^ method using the geometric mean of *Tbp* and *Rs9* as housekeeping value.

### mtDNA content

C2C12 myotubes were harvested at D6 of differentiation and digested in a buffer (0.5M KCl, Tris pH 8.0 100mM, NP40 0.45%, Tween 0.45% and Proteinase K 0.2 mg/ml) at 55°C for 2 hours. After sonication by a Bioruptor device (Diagenode), DNA was purified with 25:24:1 mixture of phenol, chloroform and isoamyl alcohol (#P3803, Sigma-Aldrich) prior to overnight precipitation in ammonium acetate and ethanol. After centrifugation, the pellet was washed with ethanol and resuspended in water. Five nanograms of DNA samples were analysed by qPCR to quantify mitochondrial and nuclear DNA using primers specific for *Nd5* and *Hk2*, respectively. qPCR was performed as described above (“qPCR on skeletal muscle and C2C12 myotube samples”).

### Western blot on skeletal muscle and C2C12 myotube samples

GNM muscles of mice euthanized in the morning in fed state were ground in liquid nitrogen and proteins were extracted from the powder obtained using Mammalian protein extraction reagent (MPER, #78501, Thermo Fisher Scientific) supplemented with halt protease and halt phosphatase inhibitors (#78429 and #78426, Thermo Fisher Scientific). C2C12 myotubes were directly harvested in MPER buffer containing inhibitors. Proteins were separated by SDS-PAGE electrophoresis and immunoblotted using antibodies targeting: S616 phospho-DRP1 (ref), S637 phospho-DRP1 (ref) and total-DRP1 (ref); subunits of the mitochondrial electron transport chain complex I (ref), complex II (ref), complex II (ref), complex IV (ref) and complex V (ref); CDK4 (ref); TOMM20 (ref); PKA-substrate proteins (ref); ESRRA (ref); PPARGC1A (ref); T172 phospho-AMPK (ref) and total-AMPK (ref); P-Rb (ref). After incubation with corresponding HRP-linked secondary antibodies, luminescence intensity was captured by a Fusion FX imager (Vilber) and quantified by densitometry using Fiji (version 2.3.0). Expression levels of proteins of interest were normalised by total protein amount (Ghosh *et al*, 2014) (Moritz, 2017) using a Pierce reversible protein stain kit (#24580, Thermo Fisher Scientific). Protein carbonylation was assessed with an Oxyblot protein oxidation detection kit (#S7150, Merck Millipore) following the manufacturer’s instructions. Briefly, proteins were extracted from GNM samples ground in liquid nitrogen with MPER buffer supplemented with protease and phosphatase inhibitors (as described above) supplemented with 50mM DTT. Carbonyl groups were then derivatized to 2,4-dinitrophenylhydrazone (DNP) and samples were separated by SDS-PAGE followed by immunoblotting with an antibody targeting the DNP moiety of the proteins. Image capture and quantification were carried out as described above for western blots.

### Statistical analysis

Statistical analyses were carried out using the Prism 9.2 software (GraphPad Software). Normal distribution was assessed by a D’Agostino-Pearson omnibus K2 test and further analyses using a Student t test (normal distribution) or a Mann-Whitney U test. When 2-way ANOVA was used, p value of each variable effect was indicated on the graph and post-hoc Fischer’s LSD tests were performed to evaluate group-to-group differences. Statistical significance was set at p value < 0.05.

## Figure legends

**Figure Supp 1:**
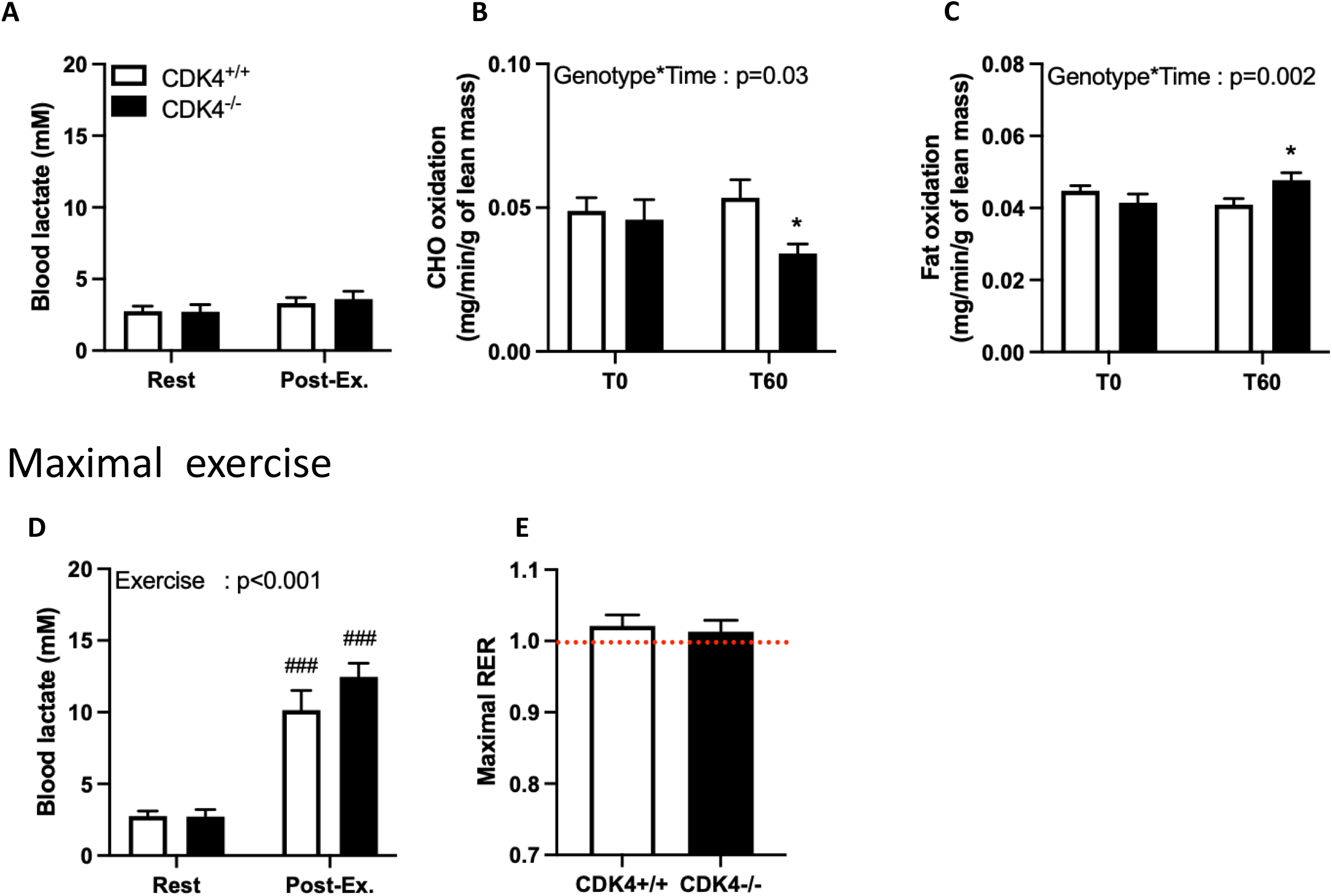
Endurance and fat oxidation during exercise are increased in *cdk4*^*-/-*^ mice. *Cdk4*^*+/+*^ and *cdk4*^*-/-*^ male mice were tested for exercise capacity between 20 and 25 weeks of age. (A) Blood lactate was measured at rest and after a 1-hour low-intensity exercise (Post-Ex) (n=11-13). (B-C) Carbohydrates (CHO) (B) and fat oxidation (C) were calculated from indirect calorimetry values at rest and after a 1-hour low-intensity exercise (Post-Ex) (n=11-13). (D) Blood lactate was measured at rest and after a graded exercise (n=11-13). (E) Maximal RER reached after a graded exercise. Empty bars: *cdk4*^*+/+*^; full bars: *cdk4*^*-/-*^. *: p<0.05 for *cdk4*^*-/-*^ vs. *cdk4*^*+/+*^. ###: p<0.001 for post- exercise vs. rest conditions.

**Figure Supp 2:**
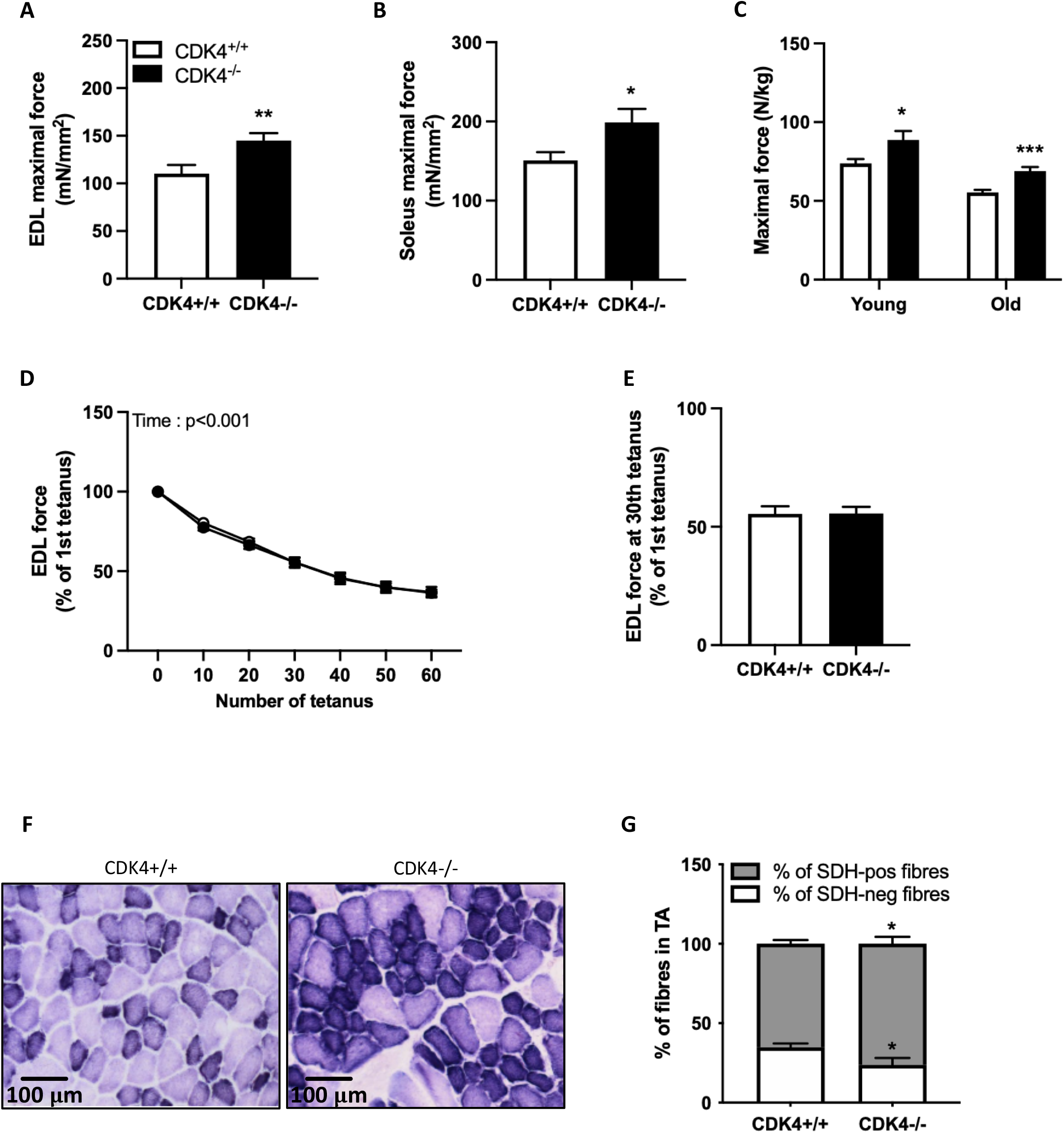
*Cdk4*^*-/-*^ mice display a skeletal muscle oxidative fibre type. *Cdk4*^*+/+*^ and *cdk4*^*-/-*^ male mice were euthanised at the age of 25 weeks and skeletal muscles were isolated. (A-B) Maximal force of extensor digitorum longus (EDL, A) and soleus (B) muscles (n=16). (C) Maximal force of mice measured in vivo during a 4-paw grip test in young (15 weeks) and old (45 weeks) mice (n=12-25). (D-E) EDL muscle fatigability was assessed by measuring maximal force production along the course of repeated tetanic contractions (n=14-15). (F) Transversal slices of tibialis anterior (TA) muscles were immunostained to visualize laminin (green) and myosin heavy chains type I (blue), type IIa (red) and type IIx or IIb (orange) (n=5). (G) Proportion of TA fibers expressing myosin heavy chain-type I, type IIa, type IIx or type IIb (n=5). (H) Transversal slices of TA were used to measure succinate dehydrogenase (SDH) activity, a marker of mitochondrial oxidative capacity. (I) Proportion of TA muscle fibers stained negatively (SDH-neg) or positively (SDH-pos) for SDH activity (n=5). Empty bars: *cdk4*^*+/+*^; full bars: *cdk4*^*-/-*^. *: p<0.05, **: p<0.01 and ***: p<0.001 for *cdk4*^*-/-*^ vs. *cdk4*^*+/+*^.

**Figure Supp 3:**
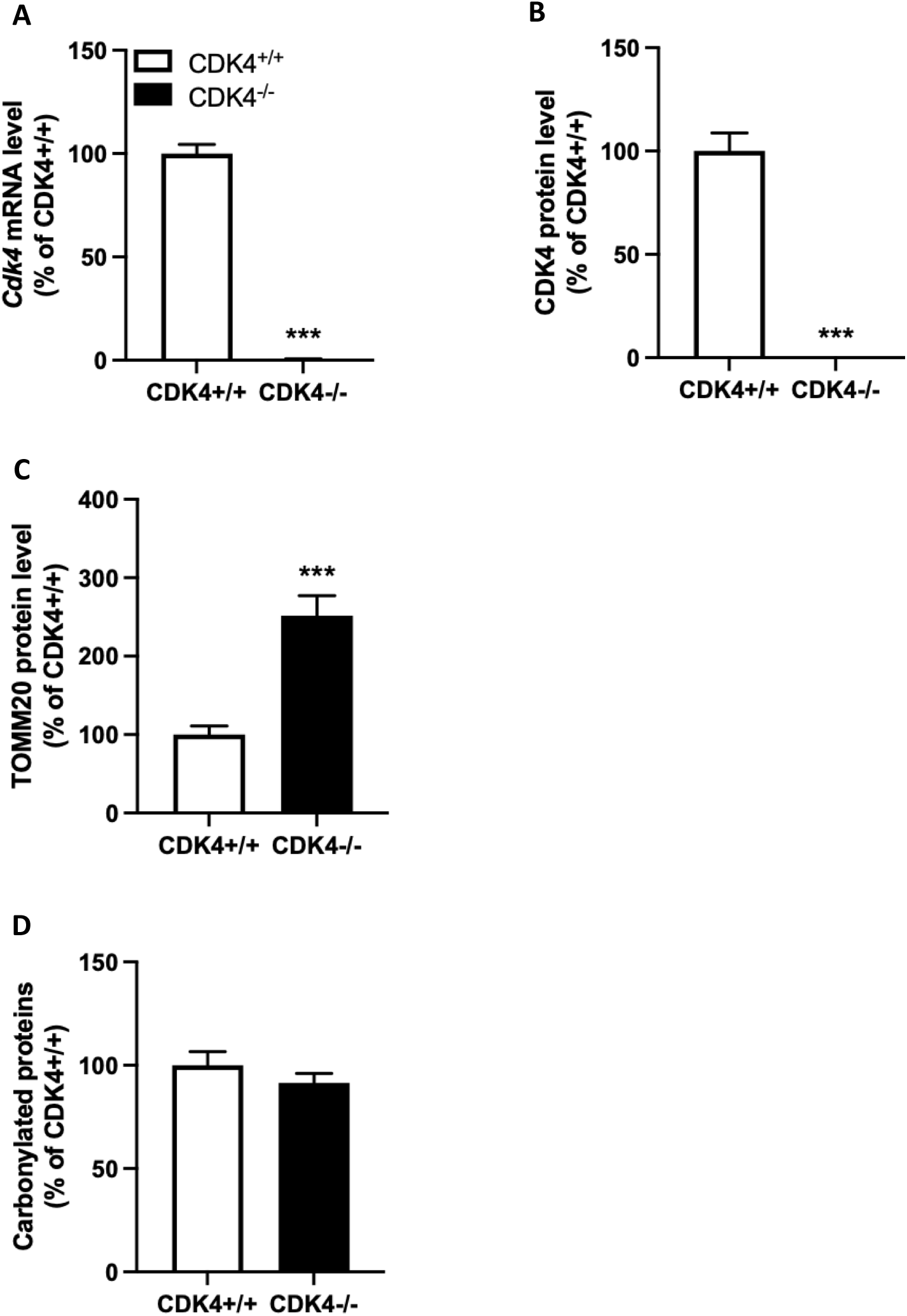
*Cdk4*^*-/-*^ mice have increased muscle mitochondrial content and oxidative capacity. *Cdk4*^*+/+*^ and CDK4-/- male mice were euthanised at the age of 25 weeks. (A-B) Gastrocnemius muscle (GNM) CDK4 mRNA (A, n=9-13) and protein (B, n=6) levels were measured by qPCR and western blot, respectively. (C) GNM TOMM20 protein levels were measured by western blot (n=6). (D) GNM content of oxidised proteins was measured by western blot using an Oxyblot kit (n=10-13). Empty bars: *cdk4*^*+/+*^; full bars: *cdk4*^*-/-*^. ***: p<0.001 for *cdk4*^*-/-*^ vs. *cdk4*^*+/+*^.

**Figure Supp 4:**
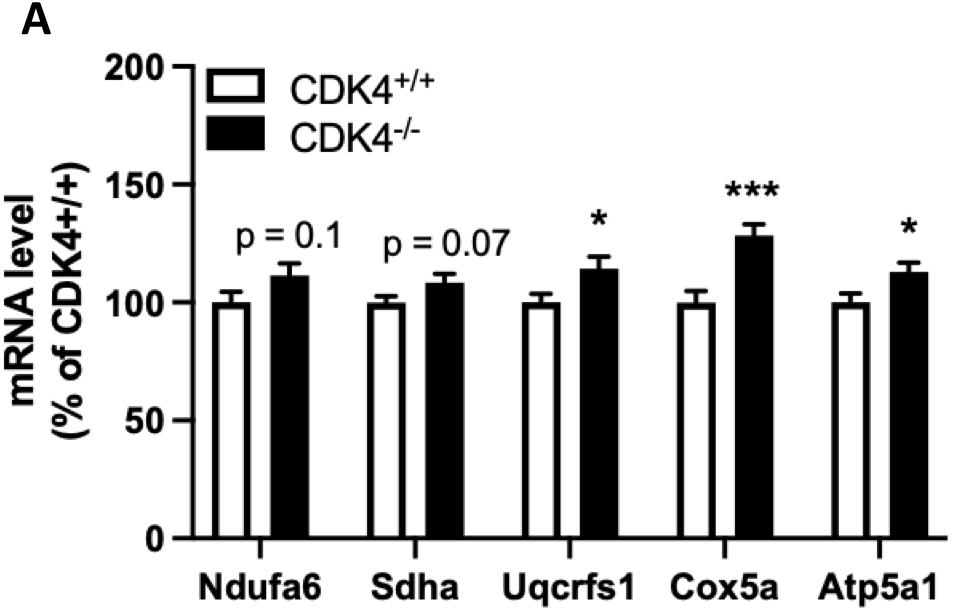
*Cdk4*^*-/-*^ skeletal muscles display an ERR-dependent transcriptomic signature. *Cdk4*^*+/+*^ and CDK4-/- male mice were euthanised at the age of 25 weeks. mRNA levels of genes coding for subunits of the mitochondrial electron transport chain (A) and of estrogen-related receptor gamma (Esrrg) (B) were measured by qPCR (n=9-13). Empty bars: *cdk4*^*+/+*^; full bars: *cdk4*^*-/-*^. *: p<0.05 and ***: p<0.001 for *cdk4*^*-/-*^ vs. *cdk4*^*+/+*^.

**Figure Supp 5:**
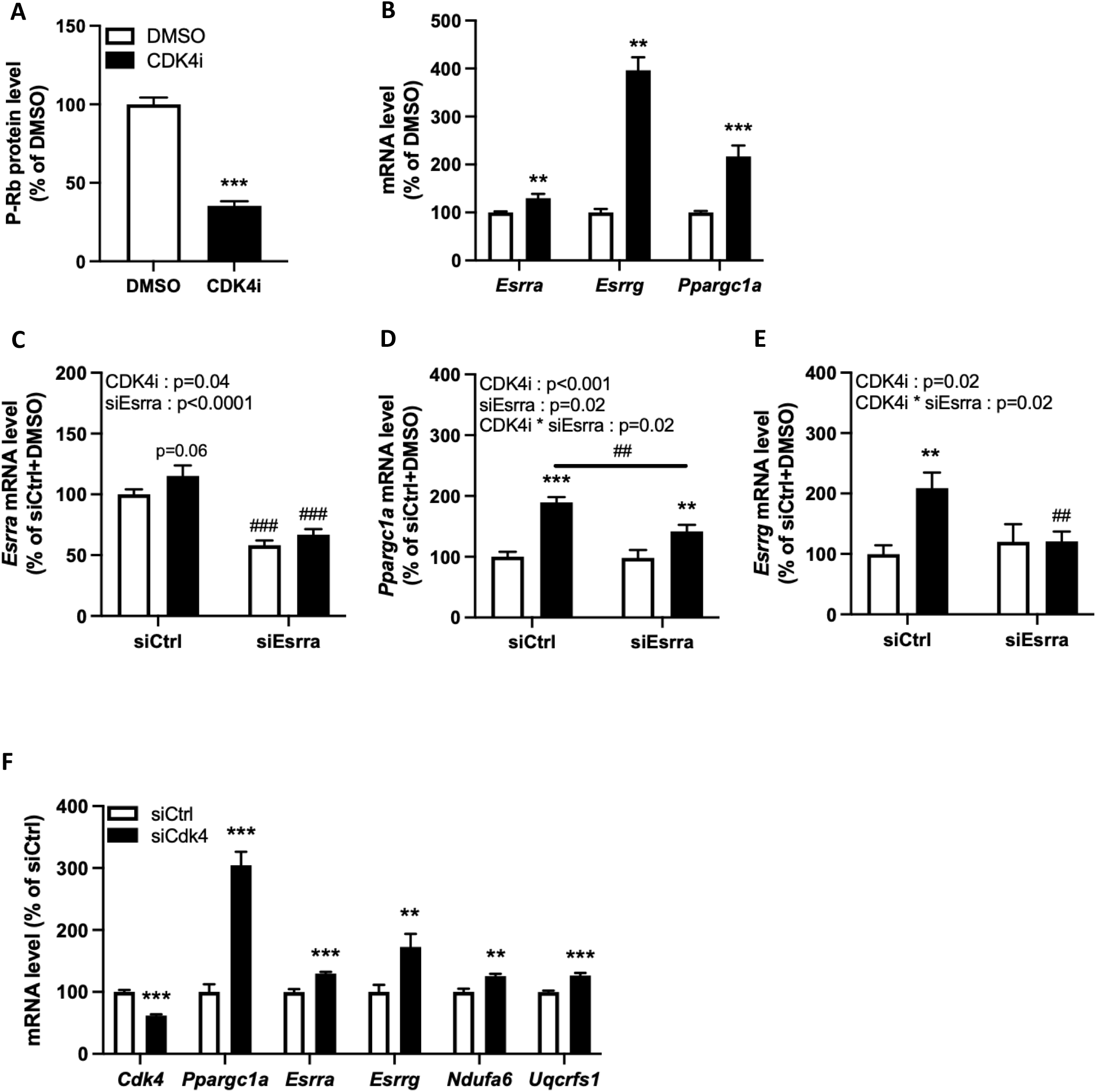
CDK4 inhibition in C2C12 myotubes stimulates ERR transcriptional activity. C2C12 myotubes were treated for 48h with abemaciclib, a CDK4/6 inhibitor (CDK4i). (A) Rb phosphorylation was measured by western blot in C2C12 myotubes treated for 48h with DMSO or abemaciclib (CDK4i) (n=8). (B) Esrra, Esrrg and Ppargc1a mRNA levels were measured by qPCR in C2C12 myotubes treated for 48h with DMSO or CDK4i (n=8). (C-E) mRNA levels were measured by qPCR in C2C12 myotubes transfected with a control siRNA (siCtrl) or a siRNA targeting Esrra (siEsrra) and then treated for 48h with DMSO or CDK4i (n=9). (F) mRNA levels were measured by qPCR in C2C12 myotubes transfected with a control siRNA (siCtrl) or a siRNA targeting Cdk4 (siCDK4) (n=9).

**Figure Supp 6:**
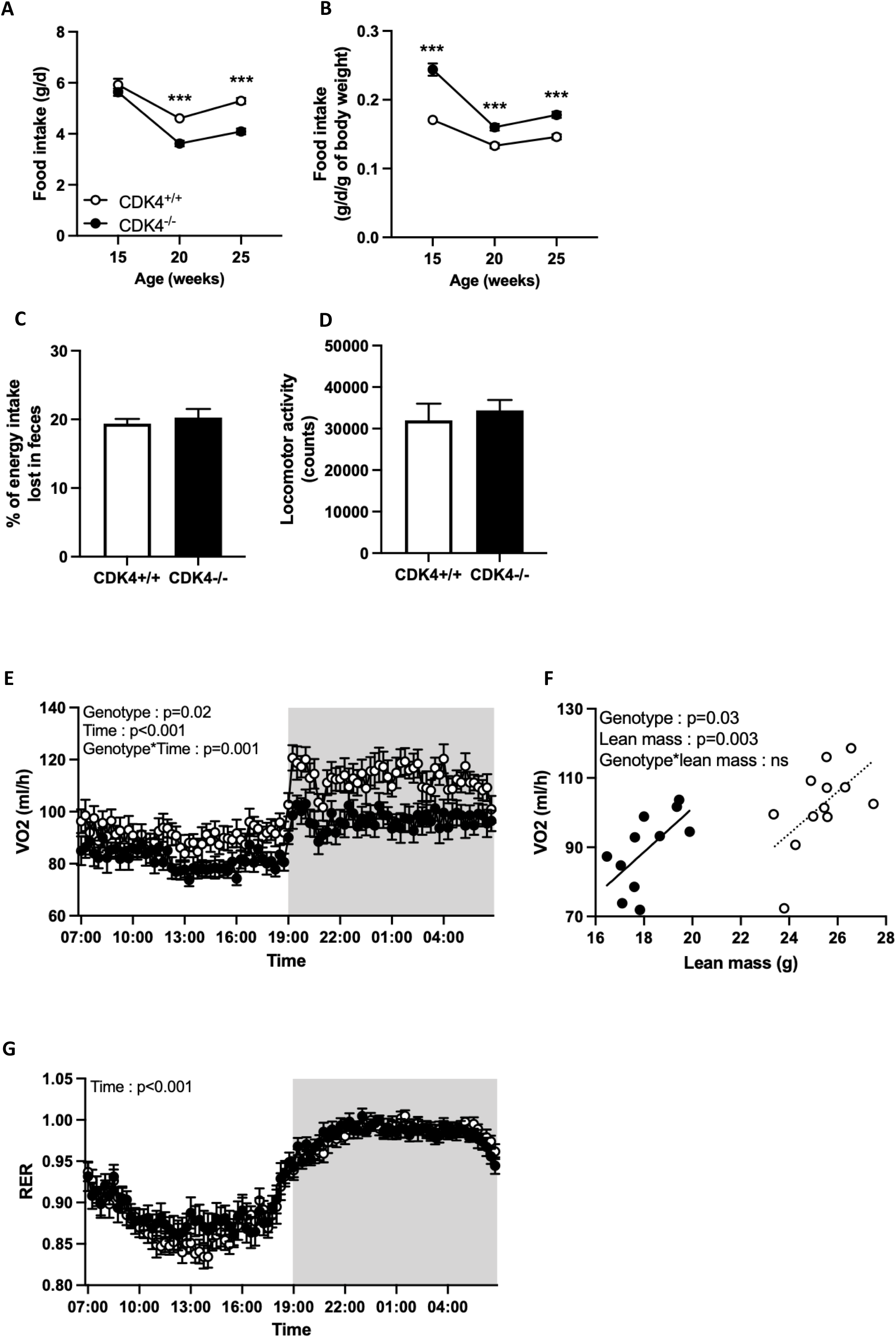
*Cdk4*^*-/-*^ mice have decreased body weight and increased metabolic rate. *Cdk4*^*+/+*^ and CDK4-/- male mice were followed from 15 to 25 weeks of age. (A) Food intake expressed in g/day and (B) in g/day/g of body weight (n=10-12). (C) Percentage of energy intake lost in feces (n=6-7). (D) Spontaneous locomotor activity measured during indirect calorimetry experiments (n=11-12). (E) O2 consumption measured by indirect calorimetry during the light (no background) and dark (grey background) phases over a 2-day period (n=11-12). (F) Average daily O^2^ consumption values were analyzed with the free online CalR application using lean mass as a covariable (n=11-12). (G) Respiratory exchange ratio (RER) calculated from indirect calorimetry measurements during the light (no background) and dark (grey background) phases over a 2-day period (n=11-12). Empty circles and bars: *cdk4*^*+/+*^; full circles and bars: *cdk4*^*-/-*^. ***: p<0.001 for *cdk4*^*-/-*^ vs. *cdk4*^*+/+*^.

## References

Achten J, Jeukendrup AE (2004) Optimizing fat oxidation through exercise and diet. Nutrition 20: 716–727

Audet-Walsh E, Giguere V (2015) The multiple universes of estrogen-related receptor alpha and gamma in metabolic control and related diseases. Acta Pharmacol Sin 36: 51–61

Ayachi M, Niel R, Momken I, Billat VL, Mille-Hamard L (2016) Validation of a Ramp Running Protocol for Determination of the True VO2max in Mice. Front Physiol 7: 372

Bosma M, Kersten S, Hesselink MK, Schrauwen P (2012) Re-evaluating lipotoxic triggers in skeletal muscle: relating intramyocellular lipid metabolism to insulin sensitivity. Prog Lipid Res 51: 36–49

Brooks SV, Faulkner JA (1988) Contractile properties of skeletal muscles from young, adult and aged mice. J Physiol 404: 71–82

Castillo-Armengol J, Barquissau V, Geller S, Ji H, Severi I, Venema W, Fenandez EA, Moret C, Huber K, Leal-Esteban LC et al (2020) Hypothalamic CDK4 regulates thermogenesis by modulating sympathetic innervation of adipose tissues. EMBO Rep 21: e49807

Fan W, Evans R (2015) PPARs and ERRs: molecular mediators of mitochondrial metabolism. Curr Opin Cell Biol 33: 49–54

George MA, Qureshi S, Omene C, Toppmeyer DL, Ganesan S (2021) Clinical and Pharmacologic Differences of CDK4/6 Inhibitors in Breast Cancer. Front Oncol 11: 693104

Ghosh R, Gilda JE, Gomes AV (2014) The necessity of and strategies for improving confidence in the accuracy of western blots. Expert Rev Proteomics 11: 549–560

Giannattasio S, Giacovazzo G, Bonato A, Caruso C, Luvisetto S, Coccurello R, Caruso M (2018) Lack of cyclin D3 induces skeletal muscle fiber-type shifting, increased endurance performance and hypermetabolism. Sci Rep 8: 12792

Giguere V (2008) Transcriptional control of energy homeostasis by the estrogen-related receptors. Endocr Rev 29: 677–696

Gomes LC, Di Benedetto G, Scorrano L (2011) During autophagy mitochondria elongate, are spared from degradation and sustain cell viability. Nat Cell Biol 13: 589–598

Goto M, Terada S, Kato M, Katoh M, Yokozeki T, Tabata I, Shimokawa T (2000) cDNA Cloning and mRNA analysis of PGC-1 in epitrochlearis muscle in swimming-exercised rats. Biochem Biophys Res Commun 274: 350–354

Hafner M, Mills CE, Subramanian K, Chen C, Chung M, Boswell SA, Everley RA, Liu C, Walmsley CS, Juric D et al (2019) Multiomics Profiling Establishes the Polypharmacology of FDA-Approved CDK4/6 Inhibitors and the Potential for Differential Clinical Activity. Cell Chem Biol 26: 1067–1080 e1068

Handschin C, Kobayashi YM, Chin S, Seale P, Campbell KP, Spiegelman BM (2007) PGC-1alpha regulates the neuromuscular junction program and ameliorates Duchenne muscular dystrophy. Genes Dev 21: 770–783

Harmuth T, Prell-Schicker C, Weber JJ, Gellerich F, Funke C, Driessen S, Magg JCD, Krebiehl G, Wolburg H, Hayer SN et al (2018) Mitochondrial Morphology, Function and Homeostasis Are Impaired by Expression of an N-terminal Calpain Cleavage Fragment of Ataxin-3. Front Mol Neurosci 11: 368

Hood DA, Uguccioni G, Vainshtein A, D’Souza D (2011) Mechanisms of exercise-induced mitochondrial biogenesis in skeletal muscle: implications for health and disease. Compr Physiol 1: 1119–1134

Huss JM, Kopp RP, Kelly DP (2002) Peroxisome proliferator-activated receptor coactivator-1alpha (PGC-1alpha) coactivates the cardiac-enriched nuclear receptors estrogen-related receptor-alpha and -gamma. Identification of novel leucine-rich interaction motif within PGC-1alpha. J Biol Chem 277: 40265–40274

Huss JM, Torra IP, Staels B, Giguere V, Kelly DP (2004) Estrogen-related receptor alpha directs peroxisome proliferator-activated receptor alpha signaling in the transcriptional control of energy metabolism in cardiac and skeletal muscle. Mol Cell Biol 24: 9079–9091

Iqbal NJ, Lu Z, Liu SM, Schwartz GJ, Chua S, Jr., Zhu L (2018) Cyclin-dependent kinase 4 is a preclinical target for diet-induced obesity. JCI Insight 3

Iqbal NJ, Schwartz GJ, Zhao H, Zhu L, Chua S, Jr. (2021) Cyclin-dependent kinase 4/6 inhibitors require an arcuate-to-paraventricular hypothalamus melanocortin circuit to treat diet-induced obesity. Am J Physiol Endocrinol Metab 320: E467–E474

Jager S, Handschin C, St-Pierre J, Spiegelman BM (2007) AMP-activated protein kinase (AMPK) action in skeletal muscle via direct phosphorylation of PGC-1alpha. Proc Natl Acad Sci U S A 104: 12017–12022

Jahani-Asl A, Huang E, Irrcher I, Rashidian J, Ishihara N, Lagace DC, Slack RS, Park DS (2015) CDK5 phosphorylates DRP1 and drives mitochondrial defects in NMDA-induced neuronal death. Hum Mol Genet 24: 4573–4583

Jeukendrup AE, Wallis GA (2005) Measurement of substrate oxidation during exercise by means of gas exchange measurements. Int J Sports Med 26 Suppl 1: S28–37

Kamarajugadda S, Becker JR, Hanse EA, Mashek DG, Mashek MT, Hendrickson AM, Mullany LK, Albrecht JH (2016) Cyclin D1 represses peroxisome proliferator-activated receptor alpha and inhibits fatty acid oxidation. Oncotarget 7: 47674–47686

Klein S, Coyle EF, Wolfe RR (1994) Fat metabolism during low-intensity exercise in endurance-trained and untrained men. Am J Physiol 267: E934–940

Koopman WJ, Visch HJ, Verkaart S, van den Heuvel LW, Smeitink JA, Willems PH (2005) Mitochondrial network complexity and pathological decrease in complex I activity are tightly correlated in isolated human complex I deficiency. Am J Physiol Cell Physiol 289: C881–890

Kremer JR, Mastronarde DN, McIntosh JR (1996) Computer visualization of three-dimensional image data using IMOD. J Struct Biol 116: 71–76

Lagarrigue S, Lopez-Mejia IC, Denechaud PD, Escote X, Castillo-Armengol J, Jimenez V, Chavey C, Giralt A, Lai Q, Zhang L et al (2016) CDK4 is an essential insulin effector in adipocytes. J Clin Invest 126: 335–348

Lee Y, Dominy JE, Choi YJ, Jurczak M, Tolliday N, Camporez JP, Chim H, Lim JH, Ruan HB, Yang X et al (2014) Cyclin D1-Cdk4 controls glucose metabolism independently of cell cycle progression. Nature 510: 547–551

Lin J, Wu H, Tarr PT, Zhang CY, Wu Z, Boss O, Michael LF, Puigserver P, Isotani E, Olson EN et al (2002) Transcriptional co-activator PGC-1 alpha drives the formation of slow-twitch muscle fibres. Nature 418: 797–801

Loomba R, Friedman SL, Shulman GI (2021) Mechanisms and disease consequences of nonalcoholic fatty liver disease. Cell 184: 2537–2564

Lopez-Mejia IC, Lagarrigue S, Giralt A, Martinez-Carreres L, Zanou N, Denechaud PD, Castillo-Armengol J, Chavey C, Orpinell M, Delacuisine B et al (2017) CDK4 Phosphorylates AMPKalpha2 to Inhibit Its Activity and Repress Fatty Acid Oxidation. Mol Cell 68: 336–349 e336

Luan P, D’Amico D, Andreux PA, Laurila PP, Wohlwend M, Li H, Imamura de Lima T, Place N, Rinsch C, Zanou N et al (2021) Urolithin A improves muscle function by inducing mitophagy in muscular dystrophy. Sci Transl Med 13

Lundby C, Jacobs RA (2016) Adaptations of skeletal muscle mitochondria to exercise training. Exp Physiol 101: 17–22

Lundsgaard AM, Fritzen AM, Kiens B (2018) Molecular Regulation of Fatty Acid Oxidation in Skeletal Muscle during Aerobic Exercise. Trends Endocrinol Metab 29: 18–30

Malumbres M (2014) Cyclin-dependent kinases. Genome Biol 15: 122

Malumbres M, Barbacid M (2009) Cell cycle, CDKs and cancer: a changing paradigm. Nat Rev Cancer 9: 153–166

Malumbres M, Sotillo R, Santamaria D, Galan J, Cerezo A, Ortega S, Dubus P, Barbacid M (2004) Mammalian cells cycle without the D-type cyclin-dependent kinases Cdk4 and Cdk6. Cell 118: 493–504

Martin J, Hunt SL, Dubus P, Sotillo R, Nehme-Pelluard F, Magnuson MA, Parlow AF, Malumbres M, Ortega S, Barbacid M (2003) Genetic rescue of Cdk4 null mice restores pancreatic beta-cell proliferation but not homeostatic cell number. Oncogene 22: 5261–5269

Mina AI, LeClair RA, LeClair KB, Cohen DE, Lantier L, Banks AS (2018) CalR: A Web-Based Analysis Tool for Indirect Calorimetry Experiments. Cell Metab 28: 656–666 e651

Moritz CP (2017) Tubulin or Not Tubulin: Heading Toward Total Protein Staining as Loading Control in Western Blots. Proteomics 17

Muller MJ, Geisler C, Hubers M, Pourhassan M, Braun W, Bosy-Westphal A (2018) Normalizing resting energy expenditure across the life course in humans: challenges and hopes. Eur J Clin Nutr 72: 628–637

Narkar VA, Downes M, Yu RT, Embler E, Wang YX, Banayo E, Mihaylova MM, Nelson MC, Zou Y, Juguilon H et al (2008) AMPK and PPARdelta agonists are exercise mimetics. Cell 134: 405–415

Puigserver P, Wu Z, Park CW, Graves R, Wright M, Spiegelman BM (1998) A cold-inducible coactivator of nuclear receptors linked to adaptive thermogenesis. Cell 92: 829–839

Purdom T, Kravitz L, Dokladny K, Mermier C (2018) Understanding the factors that effect maximal fat oxidation. J Int Soc Sports Nutr 15: 3

Rane SG, Dubus P, Mettus RV, Galbreath EJ, Boden G, Reddy EP, Barbacid M (1999) Loss of Cdk4 expression causes insulin-deficient diabetes and Cdk4 activation results in beta-islet cell hyperplasia. Nat Genet 22: 44–52

Rangwala SM, Wang X, Calvo JA, Lindsley L, Zhang Y, Deyneko G, Beaulieu V, Gao J, Turner G, Markovits J (2010) Estrogen-related receptor gamma is a key regulator of muscle mitochondrial activity and oxidative capacity. J Biol Chem 285: 22619–22629

Scarpulla RC (2011) Metabolic control of mitochondrial biogenesis through the PGC-1 family regulatory network. Biochim Biophys Acta 1813: 1269–1278

Scharhag-Rosenberger F, Meyer T, Walitzek S, Kindermann W (2010) Effects of one year aerobic endurance training on resting metabolic rate and exercise fat oxidation in previously untrained men and women. Metabolic endurance training adaptations. Int J Sports Med 31: 498–504

Schreiber SN, Knutti D, Brogli K, Uhlmann T, Kralli A (2003) The transcriptional coactivator PGC-1 regulates the expression and activity of the orphan nuclear receptor estrogen-related receptor alpha (ERRalpha). J Biol Chem 278: 9013–9018

Selsby JT, Morine KJ, Pendrak K, Barton ER, Sweeney HL (2012) Rescue of dystrophic skeletal muscle by PGC-1alpha involves a fast to slow fiber type shift in the mdx mouse. PLoS One 7: e30063

Speakman JR (2013) Measuring energy metabolism in the mouse - theoretical, practical, and analytical considerations. Front Physiol 4: 34

Taguchi N, Ishihara N, Jofuku A, Oka T, Mihara K (2007) Mitotic phosphorylation of dynamin-related GTPase Drp1 participates in mitochondrial fission. J Biol Chem 282: 11521–11529

Tanaka T, Nishimura A, Nishiyama K, Goto T, Numaga-Tomita T, Nishida M (2020) Mitochondrial dynamics in exercise physiology. Pflugers Arch 472: 137–153

Timpani CA, Hayes A, Rybalka E (2015) Revisiting the dystrophin-ATP connection: How half a century of research still implicates mitochondrial dysfunction in Duchenne Muscular Dystrophy aetiology. Med Hypotheses 85: 1021–1033

Tschop MH, Speakman JR, Arch JR, Auwerx J, Bruning JC, Chan L, Eckel RH, Farese RV, Jr., Galgani JE, Hambly C et al (2011) A guide to analysis of mouse energy metabolism. Nat Methods 9: 57–63

Wu H, Kanatous SB, Thurmond FA, Gallardo T, Isotani E, Bassel-Duby R, Williams RS (2002) Regulation of mitochondrial biogenesis in skeletal muscle by CaMK. Science 296: 349–352

Zanou N, Shapovalov G, Louis M, Tajeddine N, Gallo C, Van Schoor M, Anguish I, Cao ML, Schakman O, Dietrich A et al (2010) Role of TRPC1 channel in skeletal muscle function. Am J Physiol Cell Physiol 298: C149–162

Zong H, Ren JM, Young LH, Pypaert M, Mu J, Birnbaum MJ, Shulman GI (2002) AMP kinase is required for mitochondrial biogenesis in skeletal muscle in response to chronic energy deprivation. Proc Natl Acad Sci U S A 99: 15983–15987

